# *De Novo* assembly of the goldfish (*Carassius auratus*) genome and the evolution of genes after whole genome duplication

**DOI:** 10.1101/373431

**Authors:** Zelin Chen, Yoshihiro Omori, Sergey Koren, Takuya Shirokiya, Takuo Kuroda, Atsushi Miyamoto, Hironori Wada, Asao Fujiyama, Atsushi Toyoda, Suiyuan Zhang, Tyra G. Wolfsberg, Koichi Kawakami, Adam M. Phillippy, NISC Comparative Sequencing Program, James C. Mullikin, Shawn M. Burgess

## Abstract

For over a thousand years throughout Asia, the common goldfish (*Carassius auratus*) was raised for both food and as an ornamental pet. Selective breeding over more than 500 years has created a wide array of body and pigmentation variation particularly valued by ornamental fish enthusiasts. As a very close relative of the common carp (*Cyprinus carpio*), goldfish shares the recent genome duplication that occurred approximately 14-16 million years ago (mya) in their common ancestor. The combination of centuries of breeding and a wide array of interesting body morphologies is an exciting opportunity to link genotype to phenotype as well as understanding the dynamics of genome evolution and speciation. Here we generated a high-quality draft sequence of a “Wakin” goldfish using 71X PacBio long-reads. We identified 70,324 coding genes and more than 11,000 non-coding transcripts. We found that the two sub-genomes in goldfish retained extensive synteny and collinearity between goldfish and zebrafish. However, “ohnologous” genes were lost quickly after the carp whole-genome duplication, and the expression of 30% of the retained duplicated gene diverged significantly across seven tissues sampled. Loss of sequence identity and/or exons determined the divergence of the expression across all tissues, while loss of conserved, non-coding elements determined expression variance between different tissues. This draft assembly also provides an important resource for comparative genomics with the very commonly used zebrafish model (*Danio rerio*), and for understanding the underlying genetic causes of goldfish variants.

## Introduction

The best estimate based on mitochondrial DNA analysis from domesticated and wild-caught goldfish is that domesticated goldfish were derived from fish in southern Asia, possibly from the lower Yangtze River ^1^. More than one thousand years of ornamental breeding history has generated more than 300 goldfish variants in body shape, fin configuration, eye style and coloration ^2^, which makes goldfish and excellent genetic model system for understanding the evolution of body shape ^2^. In addition, goldfish have long been used in research to study a wide array of biological processes such as pigmentation ^3,4^, disease and environment ^5,6^, behavior ^7^, physiology ^8^, neurobiology ^9,10^, reproduction and growth ^11^, and neuroendocrine signaling ^12^.

Like the closely related common carp, goldfish experienced the same whole-genome duplication event (WGD) ≈8-12 million years ago (Mya), which is believed to have been an allotetraploidy event (i.e. the fusion without chromosome loss of two closely related species) ^13^ (figure 1c). This fusion occurred after divergence from grass carp (*Ctenopharyngodon idella*), but before goldfish diverged from the common carp. This event is quite recent compared to other animal WGD events like the one that occurred in teleosts (320-350 Mya) ^14^, in the Salmoniformes like salmon (50-80 Mya) ^15^, and the allotetraploid event of *Xenopus laevis* (17–18 Mya) ^16^, and we now have two different species that resulted from the same genome duplication event with near-complete genome sequences. Thus, comparing how the goldfish genome has diverged from the common carp provides an excellent opportunity to study how genomes change during the course of speciation. In addition, the relative evolutionary proximity of goldfish and carp to the commonly used model organism zebrafish, provides new reference sequences for identifying conserved elements involved in gene regulation (conserved non-coding elements or CNE’s) ^17,18^, at sensitivities not available from comparing much more distantly related genomes.

**Fig 1.**
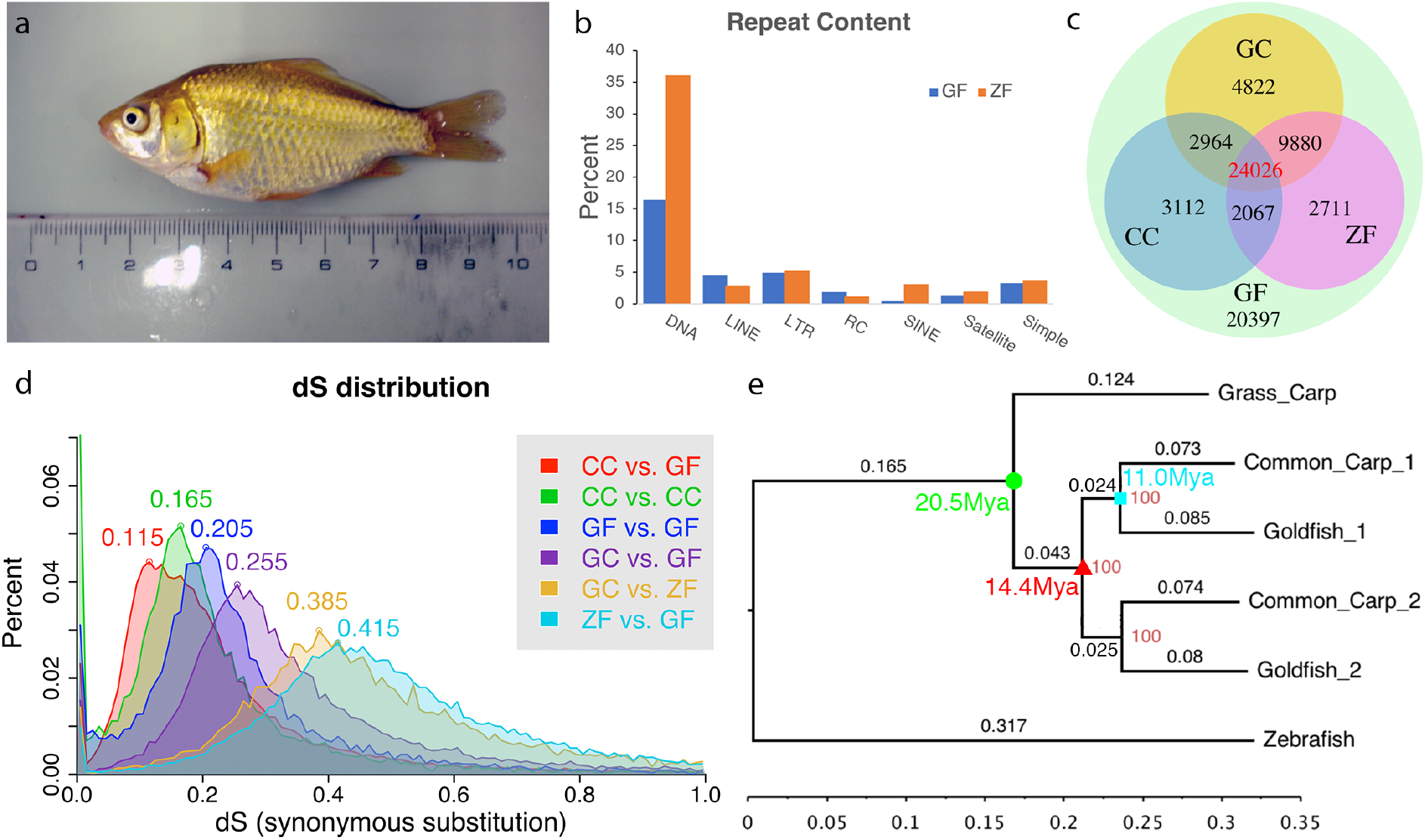
(a) The gynogenetic goldfish used for sequencing before sacrifice. (b) Transposable elements distribution for goldfish and zebrafish. (c) Distribution of orthologous/ohnologous gene pairs by synonymous substitution among four species: zebrafish, grass carp, common carp and goldfish. Numbers are a count of the homologous genes shared among zebrafish (ZF), common carp(CC) and goldfish (GF). (d) Rate of synonymous base changes (dS) for various species comparisons. (e) The phylogenetic tree shows the time of divergence of grass carp (GC) from goldfish and common carp (green circle), the whole genome duplication (red triangle) and divergence common carp and goldfish (cyan square). Each subgenome from the duplication was analyzed separately and are denoted with _1 or _2 for both common carp and goldfish. Diversion rates were similar for each subgenome.

Here we report a contiguous, accurate, and proximate-complete genome assembly of a common goldfish line, Wakin, and shed light on how the genome and gene expression evolved after the carp WGD. The genome represents an essential resource for the study of the greater than 300 goldfish variants and for the understanding of genome evolution in related fish species.

## Results

### Genomic assembly and annotation

The estimated size of the of goldfish genome ranges from 1.6 pg to 2.08 pg according to the Animal Genome Size Database ^19^, similar to that of the common carp (1.8pg). Using a Wakin goldfish generated by heat-shock gynogenesis ^20^ (figure 1a), we generated ~16.4M reads (71X coverage) from Pacbio SMRT cells, which were corrected and assembled into 9,415 contigs by the Canu assembler. The Canu assembly is ~1,849 Mbp with an N50 of 817 kbp. 6,937 contigs (497 Mbp) were of relative read coverage <0.6, which indicated that our sample was not fully homozygous with ~249 Mbp being heterozygous, consistent with the 25-mer spectrum from Illumina short-read sequencing (supplemental figure 1, supplemental table 2). We then made linkage groups using a published genetic map for the goldfish ^21^ in combination with the Onemap program ^22^. This chromosome-sized, final assembly (cauAur01) contained 50 large linkage groups (LGs), with total length of 1,246 Mbp linked and approximately 500 Mbp in unplaced contigs or scaffolds (for summary, see Table 1). By mapping the Illumina short reads to the carAur01 assembly, we estimated that the assembly has <1 error per 50,000 bases, and 98.5% reads were mappable (96% properly paired), indicating a highly accurate assembly.

**Table 1.** Assembly statistics

|  | Canu | Canu + Genetic Map |
| --- | --- | --- |
| longest | 12,834 kbp | 37,185 kbp |
| N10 | 3,990 kbp (n=135) | 30,202 kbp (n=10) |
| N50 | 817 kbp (n=504) | 22,763 kbp (n=14) |
| N90 | 64.6 kbp (n=3877) | 86.8 kbp (n=1506) |
| Total length | 1,849,050,767 bp | 1,820,635,051 bp |
| coverage by full-aligned pacbio reads (>=3) | 1,838,275,517 bp (99.41%) | - |
| No. linkage groups | - | 50 |
| Total length of linkage groups | - | 1,246,641,604 bp |

We sequenced one additional gynogenetic and one “wild-type” Wakin fish to ~70X coverage using Illumina short read sequencing. In aggregate, we identified 12,163,467 unique SNV and 2,316,524 deletion/insertion variants (DIV) from these fish, and estimated the polymorphism rate in goldfish was approximately 1%.

The goldfish genome showed an overall repeat content of 39.6%, which is similar to the 39.2% for common carp ^23^, higher than that for most of the sequenced teleost genomes (7.1% in Takifugu rubripes ^24^, 5.7% in Tetraodon nigroviridis ^25^, 30.68% in Oryzias latipes ^26^) but much lower than that of the zebrafish (54.3%) ^27^. The most enriched repeat classes were DNA transposons, of which hAT (3.87%), DNA (3.08%), TcMar (2.28%), and CMC (2.05%) were the top enriched superfamiles. Superfamilies LINE/L2 (2.67%), LTR/Gypsy (2.14%), RC/Helitron (1.89%), and LTR/DIRs (1.18%) were also somewhat enriched (>1%). Goldfish contains more LINEs but fewer SINE and DNA transposons than zebrafish (figure 1b and supplemental table 3). A fully implemented UCSC browser of carAur01 is available at: https://research.nhgri.nih.gov/goldfish/ (supplemental figure 2).

We sequenced and assembled total RNA from seven adult tissues (brain, gill, bone, eye heart, skeletal muscle, and fin). Maker identified 80,062 protein coding genes 9,738 genes were masked because they were duplicated in the heterozygous regions. The final assembly, carAur01, contained 70,324 unmasked gene models and 479,594 exons. The gene completeness was assessed by Benchmarking Universal Single-Copy Orthologs (BUSCO) ^28^ using the vertebrate core gene sets, resulting in 2,710 complete (90%), 157 fragmented (5%), and 156 (5%) missing BUSCOs out of 3,023 total BUSCOs (see table 2 and supplemental table 4). 58% of the BUSCO genes could be found in two complete copies. 83.11% to 96.93% of the RNA-seq reads from seven goldfish tissues could be mapped to the assembly. These assessments indicated our gene models were of very good quality and significantly more complete than that of the published common carp assembly. Based on Ensembl alignment evidence, we predicted 11,820 non-coding RNA transcripts, include 574 micro RNAs. miRBase hairpin sequence alignment identified 1,037 microRNA loci.

~50,000 coding gene had a RBH (reciprocal best hit) or second best hit to genes in zebrafish, grass carp or common carp. 24,026 genes hit to all three species (figure 1c). The spectrum of synonymous substitutions (dS) between RBH pairs showed peaks at 0.115, 0.205, 0.415 for common carp-goldfish (figure 1d, CC vs. GF), between goldfish WGD paralogs (figure 1d, GF vs. GF) and zebrafish-goldfish (figure 1d, ZF vs. GF) comparisons respectively. As expected, this indicated that the whole genome duplication event happened before the divergence of goldfish and common carp. Based on the ML phylogenetic tree and using 20.5 mya for the grass carp – common carp divergence point, we deduced the speciation time for common carp and goldfish was ~11.0 Mya and the WGD time was ~14.4 Mya (figure 1e), which is consistent with Larhammer and Risinger’s estimate ^29^, but slightly longer ago than other more recent publications’ predictions ^13,23^.

### Extensive retention of synteny and collinearity after WGD

Though goldfish diverge from zebrafish ~60 mya, the genome of goldfish retained extensive collinearity/synteny with that of zebrafish. 97.4% of RBH or second best ortholog gene pairs between goldfish and zebrafish are located in the 25 synteny triples, including one zebrafish chromosome and two corresponding goldfish LGs. No large inter-chromosome translocations were found between the 25 zebrafish chromosomes and the 50 goldfish LGs (figure 2). This is consistent with the WGD (allotetraploid) hypothesis ^13^. Alignment between zebrafish chromosome and two WGD descended goldfish LGs shows large collinear block, thought there are large intra-chromosomal rearrangements (figure 3, supplemental figure 7), which indicated that the gene order in goldfish genome remained stable after divergence from zebrafish.

**Fig 2.**
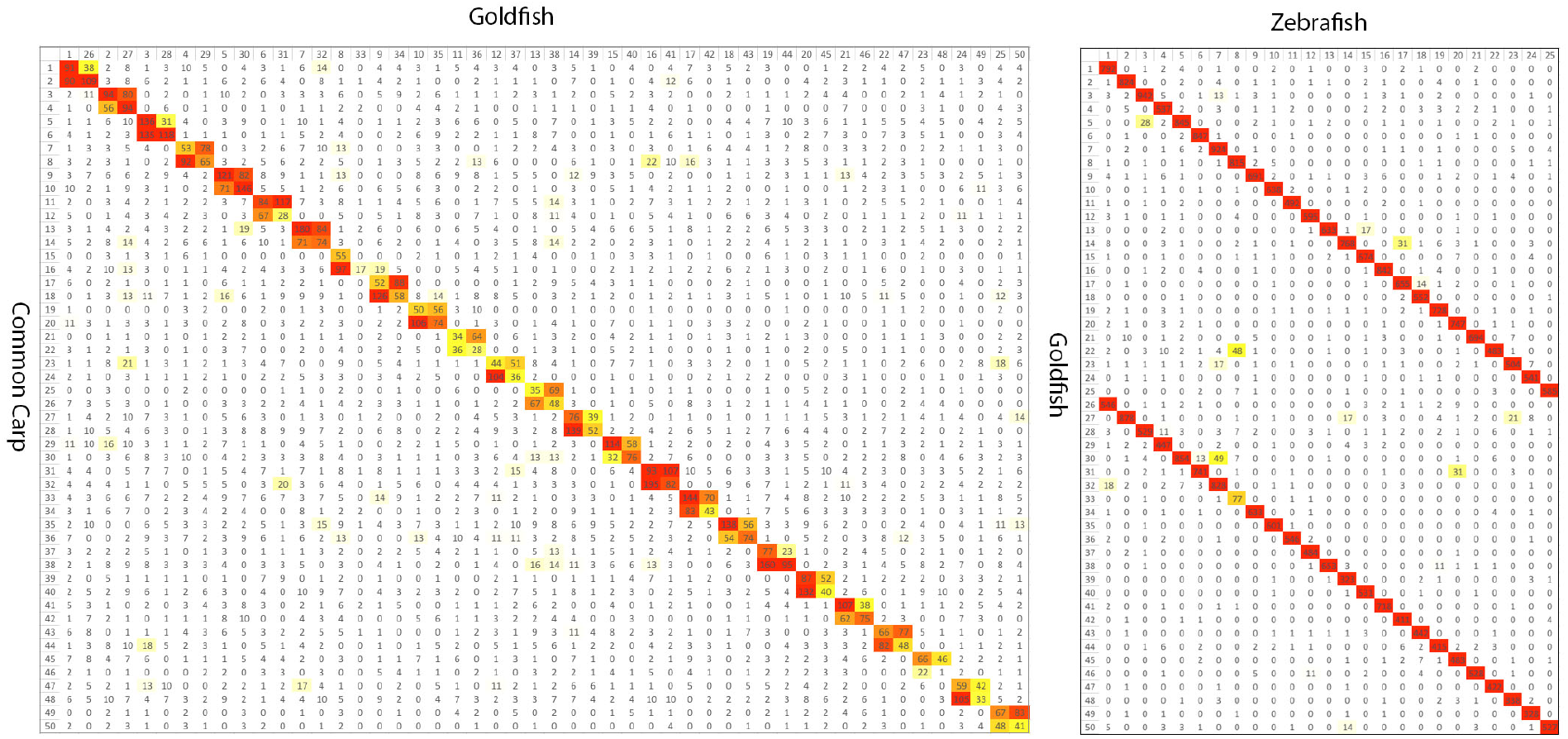
Reciprocal BLAST best gene pair counts for each pair of chromosomes. Left: goldfish and common carp. Right: goldfish and zebrafish. Color from yellow to red indicates low to high counts. Goldfish to common carp results in 50 bivalents and goldfish to zebrafish shows a clear 1:2 relationship.

**Fig 3.**
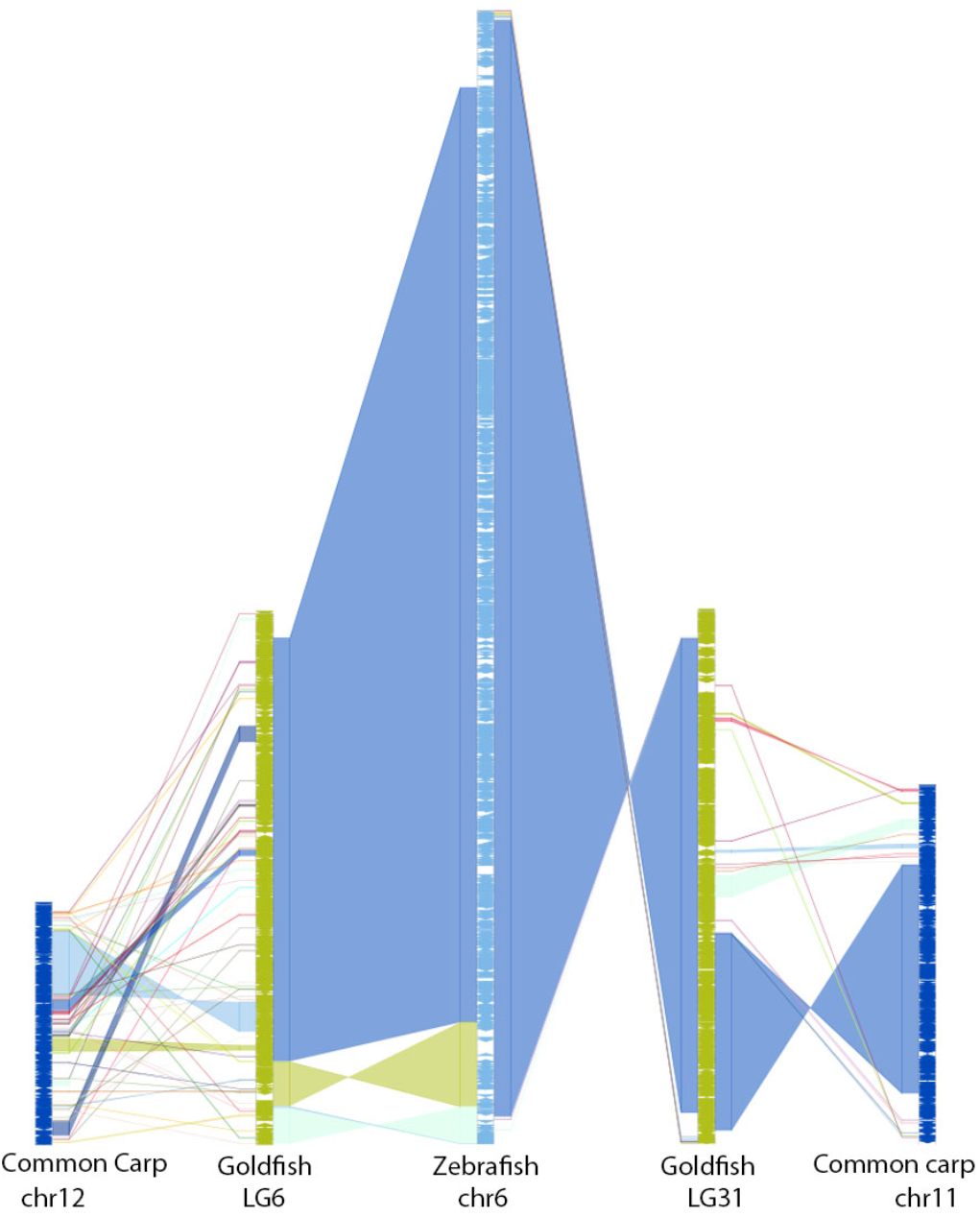
Chain alignment along zebrafish chromosome 6 and the two duplicated chromosomes from goldfish and common carp. Very large stretches of collinearity are readily visible between zebrafish and goldfish as are simple intra-chromosomal inversions. The more fragmented relationship with common carp (e.g. chr12) may be a result of a more fragmented assembly.

Only 55.3% of RBH orthologous pairs were located in the 25 LG quadruplets (2 goldfish paralog LGs and 2 common carp paralog LGs derived from the same WGD ancestral chromosome), and there are also plenty of inter-chromosomal translocations between the paralog LGs, suggesting intensive inter-chromosome translocations between common carp LGs after the WGD, especially after speciation from goldfish (figure 2). Comparisons between common carp and goldfish orthologous LGs suggested there were some small, inter-chromosome translocations though they maintained very strong colinearity (figure 3, supplemental figure 7).

### Evolution after whole genome duplication

Four available fish genomes in the *Cyprinidae* family, zebrafish, grass carp, common carp, and now goldfish, possess a very useful evolutionary relationship that allows us to directly examine the processes of gene nonfunctionalization, subfunctionalization, and neofunctionalization ^30^ over a short time (10~20 *My*) after WGD. Zebrafish is distantly and equally related to all three carps (common ancestor was ~60 mya, roughly similar to a human to mouse genomic comparison), such that the conserved sequences from zebrafish to carp are limited to exonic sequences and conserved non-coding elements (CNEs) ^17,18^ that are strongly enriched for enhancers and promoters. Common carp and goldfish speciated from grass carp ~20 Mya ^31^, the genome duplication occurred ~14 Mya and then goldfish and common carp speciated roughly 11 Mya (figure 1e). This timeline allows us to watch as duplicated genes naturally decay from the tetraploid state as was done for common carp ^32^, and the common carp, goldfish separation allows us to watch this occur twice in parallel.

#### Gene loss

We should be able to map one grass carp or zebrafish gene to two goldfish or common carp ohnologous genes. We identified 17,950 ortholog-paralog gene clusters with at least one zebrafish gene in each cluster. There are 15,011 (11,812) clusters with both paralogs retained and 2,503 (5,030) singletons in goldfish (common carp). Therefore, 14% of the duplicated gene pairs have lost one copy in goldfish while common carp appears to have had a higher rate of gene loss (28%) (supplemental figure 8). The higher loss rates in common carp may reflect the more fragmented assembly of that genome and not an actual increase in gene loss as is suggested through the lower completeness of the BUSCO genes in the common carp assembly (Table 2). Additionally, 649 (3.6%) of clusters with both ohnologs retained do not express both ohnologs in any of the seven tissues, suggesting they may be pseudogenes. In total 18% (1.3% per *My*) of WGD ancestor genes lost function in one ohnolog in goldfish during ~14 *My*, compared to 45% (0.56% per *my*) loss in salmon during 80 *My* after the salmon WGD ^15^ and the approximately 10% gene loss that occurred between zebrafish and grass carp over 60 *My*, suggesting gene loss rate increased after WGD event, which is supported by the observed faster loss (44% in 18 my or 2.4% per *My*) in *X. laevis* after the frog allotetraploid event ^16^. We then went on to ask if there were specific classes of genes that were either more or less likely than average to be lost. We examined the percentage of genes in a GO term category that were lost compared to the total percentage the category represented. Oxidoreductase activity, nuclease activity, and methyltransferase activity were much more likely than average to be lost, while protein binding and transcription factors were retained at a higher than average rate (see supplemental figures 9).

**Table 2.**
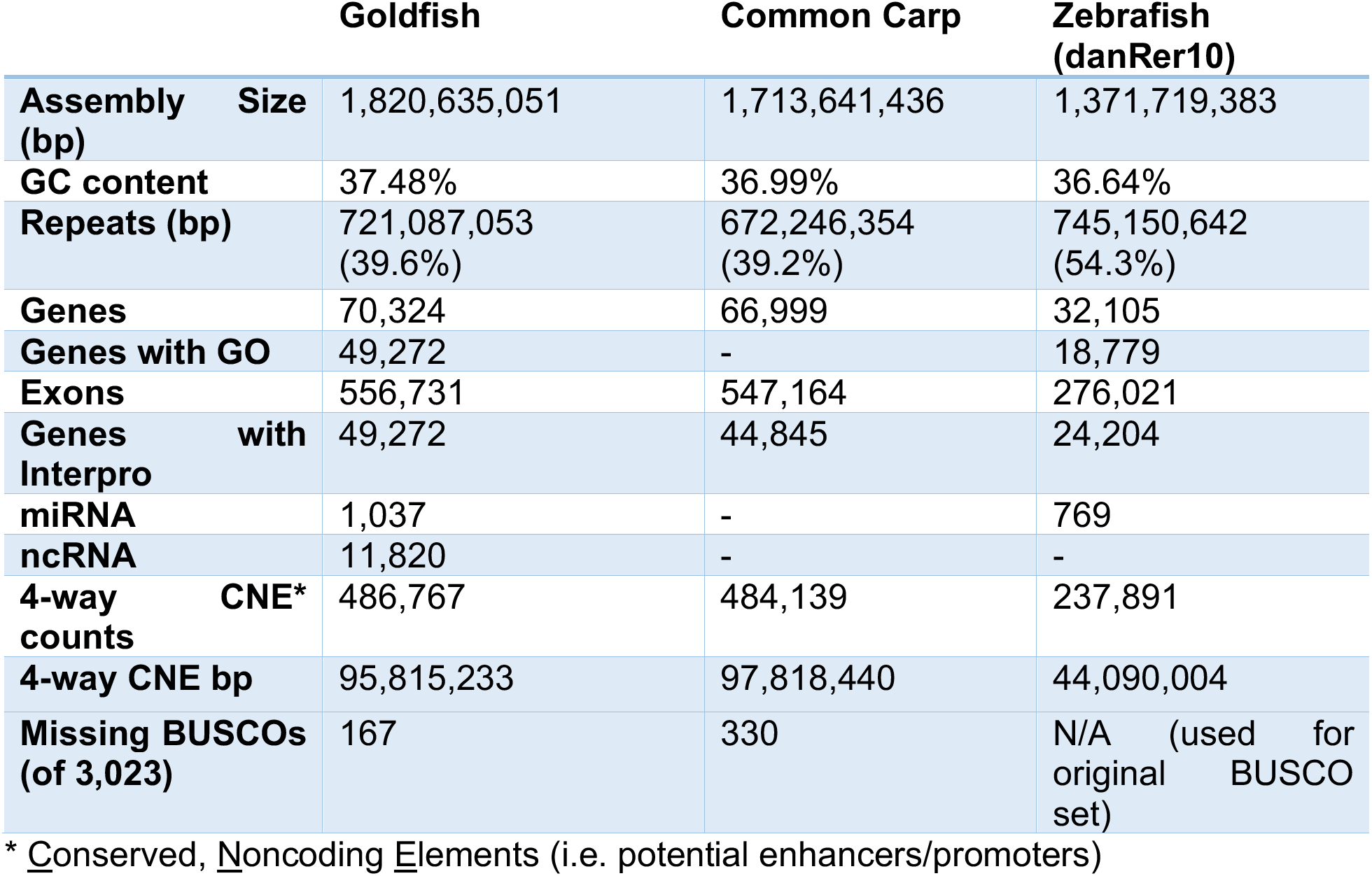
Annotations statistics

|  | **Goldfish** | **Common Carp** | **Zebrafish**<br>**(danRer10)** |
| --- | --- | --- | --- |
| **Assembly Size (bp)** | 1,820,635,051 | 1,713,641,436 | 1,371,719,383 |
| **GC content** | 37.48% | 36.99% | 36.64% |
| **Repeats (bp)** | 721,087,053<br>(39.6%) | 672,246,354<br>(39.2%) | 745,150,642<br>(54.3%) |
| **Genes** | 70,324 | 66,999 | 32,105 |
| **Genes with GO** | 49,272 | - | 18,779 |
| **Exons** | 556,731 | 547,164 | 276,021 |
| **Genes with Interpro** | 49,272 | 44,845 | 24,204 |
| **miRNA** | 1,037 | - | 769 |
| **ncRNA** | 11,820 | - | - |
| **4-way CNE\* counts** | 486,767 | 484,139 | 237,891 |
| **4-way CNE bp** | 95,815,233 | 97,818,440 | 44,090,004 |
| **Missing BUSCOs (of 3,023)** | 167 | 330 | N/A (used for original BUSCO set) |

\* Conserved, Noncoding Elements (i.e. potential enhancers/promoters)

#### CNE loss

We were able to analyze enhancer/promoter loss rates in a four-way comparison using CNE loss as the proxy for regulatory function. When we directly compared zebrafish and grass carp (using common carp or goldfish as the reference), 15,745 CNEs were not shared between them. Assuming they were lost either in zebrafish or grass carp, we estimated the lost rate was 131 CNEs per *My*. Using zebrafish as the reference, 3,611 CNEs were lost during the 40 *My* (or 90 CNEs per *My*) to grass carp. There are 329 CNEs (54 CNEs per *My*) where the two duplicated copies are missing in both goldfish and common carp. These are CNE losses that presumably happened after the split from grass carp, but before the whole genome duplication. Goldfish and common carp share 4,316 one-copy CNE losses, presumably all or most of those occurred in the 3 *My* between the genome duplication and speciation events, resulting in a rate of 1,439 per *My*. In the ~11 *My* since the common carp/goldfish split, 16,102 and 28,937 CNE paralog pairs became singleton or totally lost in goldfish and common carp respectively, or 1,463 and 2,631 CNEs per *My*. (supplemental figure 8). The above scenario indicates an accelerated CNE loss after the WGD and the effect persisted after the speciation of goldfish and common carp.

#### Divergence of gene expression

It is logical to assume that as a genome goes through the evolutionary process of re-diploidization, genes that were once duplicates of each other, will begin to diverge in location of expression or in specific function from each other. The goldfish/carp duplication event was relatively recent, which make it possible to illuminate how sequence divergence, exon loss, and CNE loss shaped the expression pattern of ohnolog genes in the ~14 *My* after the whole genome duplication. We identified 2,481 co-linear ohnolog blocks covering 1,004 Mbp of the carAur01 assembly, including 44,650 protein coding genes (6,385 singleton), 14,527 singleton exons and 8,617 singleton CNEs.

We compared the RNA expression level between 10,399 ohnolog gene pairs (20,798 genes) in the ohnolog blocks across 7 tissues. 6.2% (649) of these gene pairs contained one silenced gene (*i.e*. TPM<1 in all tissues), which may be genes that have become non-functionalized or simply not expressed in the tissues profiled. The silenced genes showed a significantly higher rate of exon loss compared to the other genes (Fisher’s exact test, p=2.2e^-16^). 2,895 (29.7%) of the remain ohnolog pairs showed divergent expression (i.e. Pearson correlation coefficient < 0.6 or Euclidean distance >=5) (figure 4a). 1,273 (13%) ohnolog pairs were tissue-specific (i.e. one gene expressed in one tissue, while the other gene silenced in the same tissue).

In order to illuminate which type of mutations contributed most to the divergence of the expression between ohnolog gene pairs, we divided these gene pairs into different groups according to their cDNA sequence identity, number of exons lost, or number of CNE lost and looked for correlations between group assignment and expression divergence. We found that in the low sequence identity groups there was greater percentage of diverged gene pairs and a lower percentage of diverged gene pairs in the high sequence identity groups (figure 4b yellow line), while the trend was reversed for less diverged gene pairs (figure 4b blue line), indicating that expression distance increased as the sequence identity decreased. There is significant increase in expression distance between the no-exon-lost group and the one-exon-lost group (one-sided Fisher exact test p=5.87e-07). The more exons were lost, the more the expression diverged (figure 4c). We did not find a significant relationship between the number of nearby CNE lost and the expression distance or correlation. However, in the ohnolog gene pairs with CNE loss but no exon loss, the tissue expression standard deviation decreased in the genes that lost CNEs (one-sided Fisher exact test p=0.008), which indicated that the loss of CNE reduced the expression variance among different tissues, rather than affecting the expression divergence between ohnolog gene pairs. *I.e*. CNE loss reduced tissue specific expression differences (figure 4d, example in supplemental figure 11)^33^.

19,500 genes (or 9,750 gene pairs, not include the silenced singletons) were classified into 20 clusters according to a plateau in their expression Euclidean distance (figure 4e and supplemental figures 12-14). Ohnologs were classified into different clusters in 62.4% of gene pairs, which decrease to 46.9% when we classified into 8 clusters (another local plateau), suggesting either a rapid expression divergence between ohnolog gene pairs in the first ~14My after the WGD event or some significant differences in gene expression that existed before the allotetraploid fusion event. Most of shared gene pairs fell within two super clusters, clusters 1-9 (figure 4e, blue curve bundles) and clusters 12-20 (figure 4e, red curve bundles). However, there are 2,508 gene pairs that are not in the same cluster within the two different super clusters. We found that there are fewer numbers of genes with lost exons or CNEs in the four most highly expressed clusters (10,11,12,15), especially in the highest expression cluster 10, in which there are no exon or CNE losses between the pairs. Similar to gene loss, genes that were more likely to maintain concordant expression were often involved in cell signaling and gene regulation (signaling molecules and transcription factors) (supplemental figure 15).

**Figure 4.**
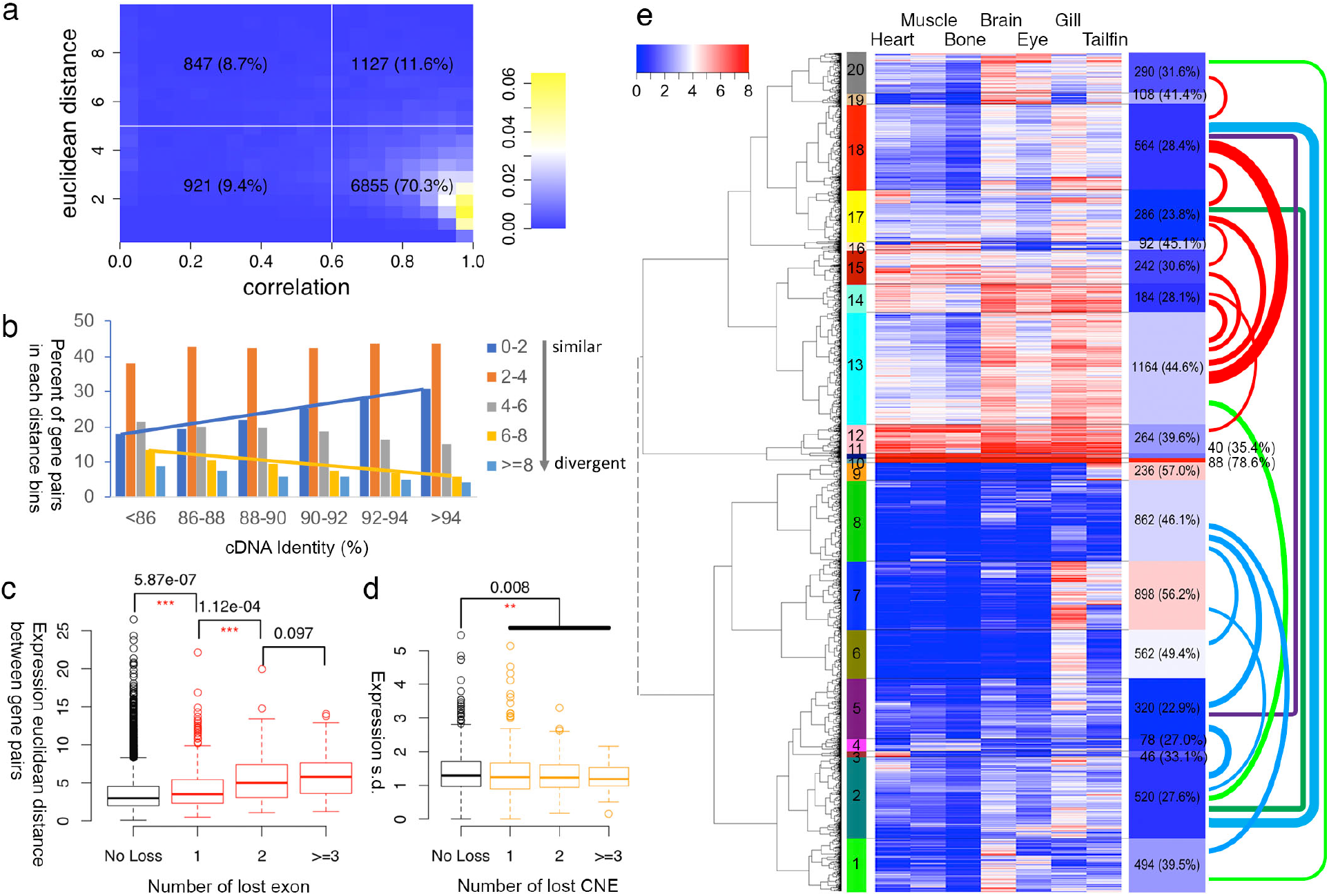
(a) Histogram of expression correlation (X-axis) and expression Euclidean distance (Y-axis) between WGD ohnolog gene pairs. Each box lists the number of ohnolog pairs (X2 for total genes) and the percentage of the total number of pairs this group represents. The majority of genes (70.3%) had a correlation of 0.6 or better. (b) expression distance distribution in different cDNA identity groups. The more closely related the cDNA sequence, the more closely correlated gene expression was. (c) Boxplot of expression distance in gene groups with different numbers of lost exons. The more exons lost the less related gene expression becomes. Asterisks mark statistically significant differences. (d) Boxplot of tissue expression standard deviation in gene groups with different numbers of CNEs lost. Similar to exons, loss of CNEs correlates with loss of concordant expression, but the effect size is smaller. Asterisks denote significant differences. (e) Gene expression clustered into 20 groups for the 19,500 ohnolog genes. Heatmap and the keys indicate the value of log_2_(TPM+1). Left color bar indicates different clusters. Right bars show the number and percentage of the gene pairs in the same cluster. Colored links indicate the number of gene pairs split between different clusters, only numbers large than 100 were plotted, thicker links indicate larger counts.

## Discussion

Steady improvements in sequencing technology and reductions in cost are improving our ability to generate high-quality genomic sequences, even in cases such as the goldfish, where the efforts are complicated by a recent whole genome duplication. Interest in the goldfish has a long history and goldfish still maintain a special position in both the scientific and ornamental fish communities. We have generated and made publicly available, a high-quality, annotated assembly of the goldfish genome. Our genomic assembly and gene annotations represents an important resource to these communities as they continue to link phenotypes to genotypes. In addition, the cluster of sequenced genomes that includes zebrafish, grass carp, common carp, and goldfish are nicely situated in their evolutionary relationship to provide further insights into the process of re-diploidization after a whole genome duplication. Comparing loss rates between that of zebrafish to grass carp and zebrafish to goldfish, despite a lower quality assembly, grass carp shows half as many gene losses as goldfish consistent with a hypothesis of accelerated gene copy loss after the whole-genome duplication. However, some functional classes of genes such as transcription factors were more likely to be preserved in two copies. Several other features of genome sequence evolution impact how gene pairs diverge in expression over time. Key factors include divergence of the primary genomic sequence through base substitution, loss of exons through deletion, and loss of conserved, non-coding elements, all of which impact gene expression in different ways. This process is one that has been proposed to be a critical evolutionary phenomenon that drives vertebrate diversity and the goldfish/carp speciation is a useful case to explore this evolutionary process.

## Acknowledgements

This work was supported by Grant–in-Aid for Scientific Research (C) (16K08583 to Y.O.) from the Japan Society for the Promotion of Science (JSPS) and NIG Collaborative Research Program (2016B5) to Y.O. This research is also funded by the Intramural Research Program of the National Human Genome Research Institute; National Institutes of Health (S.M.B.: 1ZIAHG000183; J.C.M.: 1ZIBHG000196; and A.M.P.: 1ZIAHG200398).

## Methods

### Additional methods and assembly information is included in supplementary materials

#### Preparation of genomic DNA and total RNA from goldfish

Gynogenic offspring were generated as previously described with some modifications ^20^. The Wakin goldfish eggs were treated with common carp sperm irradiated by UV-light (8000erg/mm2). After 34 min incubation at 20°C the eggs were subjected to post-fertilization heat-shock treatment at 40°C for 40 sec. After 1 min incubation at 20°C the eggs were subjected to second heat-shock treatment at 40°C for 40 sec. After heat-shock treatment the fertilized eggs were incubated at 20°C. The muscle tissue was dissected from gynogenic goldfish at 3 months of age, and high molecular weight Genomic DNA were purified using TissueLyser II (Qiagen) and Blood & Cell Culture DNA Maxi Kit (Qiagen). The molecular size of genomic DNA at the peak of 40-to 50-kb was confirmed using Pippin pulse electroporation system (NIPPON genetics). Tissues for RNA-seq were dissected from Wakin goldfish at two years of age and were stored in RNAlater (Sigma) at −80°C. Total RNA was purified using TRIzol reagent (Invitrogen) from these tissues. All procedures were approved by the Animal Experimental Committees of the Institute for Protein Research at Osaka University, and were performed in compliance with the institutional guidelines.

#### Genome Assembly

Genomic DNA from the samples described above were used to perform whole-genome shotgun sequencing on a PacBio RS II sequencer. ~16.4M Pacbio subreads (~71X) with peak length of ~8kbp were corrected and assembled into 9,415 contigs using the Canu assembler and improved the accuracy using Arrow ^34^. Total length of the assembly is 1,848 Mb and N50 reached 816.8kbp. The longest contig is 12.8Mbp. We remapped all Pacbio subreads to this assembly and found that 6,607 contigs had read coverage less than 0.6 with a total length is 596 Mbp. The reason for this appears to be the heatshock gynogenesis resulted in a meiosis II block creating heterogeneous diploid regions in approximately 22% our fish genome, as opposed to the expected mitosis I block that would have been fully homozygous. It is possible the fully homozygous fish in the heat shocked samples were not viable because of too many potentially harmful mutations in the background. The homozygous regions had 2,667 contigs (total length ~1,247Mbp) with read coverage in a range of 0.6 to 1.8. This is consent with results from our Illumina short-read sequencing which indicated about 1/4 of the genome was heterogeneous. By summing all contig length normalized by read coverage, we obtained an actual genome size of at least 1.6Gbp. To remove the alternate alleles from the primary assembly, all contigs were aligned to one another other using nucmer ^35^ and 928 contigs fully contained in other contigs were removed (when relative read coverage was <0.6 and identity was >97% to prevent WGD ohnolog removal), which was 27.3Mbp in total.

#### Linkage Group Construction

RNA-seq data from two goldfish parents and their family were download from NCBI (bioproject:PRJEB12518) ^21^. All reads were trimmed using Trimmomatic (same configuring as in Gene Annotation) and aligned to the Canu assembly using Hisat2 ^36^.

Variant calling was performed via samtools mpileup and bcftools call (parameter ‘-m’) ^37^. We identified ~5.6 M variants in total. SNPs without a matching genotype or low read depth (<4) in more than 25% of the samples, or with a missing genotype from one of the two parents were removed (other filter: bcftools filter -g 10 -Ov -i ‘TYPE=“snp” && QUAL>=10 && INFO/DP>=50’). SNPs that were homozygous in both parents or failed a Mendelian test were also removed. We also made sure two SNPs on the same contig were separated by at least 10Kbp. 14,022 SNPs were kept after filtering and used for constructing the genetic maps.

SNPs from same contigs were grouped and ordered using ‘group’ and ‘seq.order’ from R package ‘onemap’ ^22^, with LOD threshold 5.5. Contigs with two or more groups (with each >= 3 markers) were broken at the position where read depth valley and depthwas < 20 and depth was in the < 20% quantile. In total, 16 contigs were broken. Contigs were placed in each linkage group according to the ordered SNPs using chromonomer. After manual corrections, 50 long linkage groups were retained and named according their alignment to the zebrafish genome (e.g. LG1 and LG26 map to zebrafish chr1, LG2 and LG27 map to zebrafish chr2, etc.).

#### Conserved Noncoding Element Annotation

All-to-all pairwise genomic alignment was performed using lastz (--gapped -- ambiguous=n --step=10 --strand=both --masking=10 --maxwordcount=500 -- identity=70..100 --format=axt) and axtToChain for four species (goldfish, common carp, grass carp, zebrafish). Alignments in repeat regions were subtracted and transformed to maf format, splitting at gaps longer than 30bp (chainToAxt –maxGap=30, then axtToMaf-score). All the pairwise MAF files were transformed to multiple alignment MAF files using roast (P=multic). Phylogenetic model were fit for each chromosome, linkage group or scaffold using phyloFit (--tree ‘(ZF,(GC,(GF,CC)))’ --subst-mod REV --nrate 4), which was used by phastCons for computing conserve score and regions. The conserved regions out of exons (of coding or noncoding genes) were defined as conserved noncoding elements for each of the four species. DNA sequence were also extracted from these elements.

#### Data deposition

PacBio raw reads have been deposited in the SRA, Project ID: PRJNA481500. The BioSample accession is: SAMN09670328. Canu assembly deposited in GenBank under accession number QPKE00000000.

Data release date Aug 1^st^, 2018.

## Supplementary Methods and Analysis

### Goldfish Genome Homepage

https://research.nhgri.nih.gov/goldfish/

### *De novo* Assembly

#### Goldfish husbandry

Fertilized goldfish eggs were incubated at 20°C. After 3 to 5 days post-fertilization (dpf), hatched goldfish larvae were fed brine shrimp (Artemia) twice per day. The water in tanks for larvae was changed with fresh water incubated at 20°C every week. After 14 dpf, goldfish were fed pellets once per day. The water in tanks for adult goldfish was changed with fresh water every month. All procedures using goldfish were approved by the Animal Experimental Committees of the Institute for Protein Research at Osaka University (approval ID 29-03-0), and were performed according to the Guidelines for Animal Experiments of Osaka University.

#### Genome Assembly

We obtained 16,671,136 reads longer than 1kbp, containing a total of 130 Gb with an N50 length of 9,889 bases (table 1). All reads were corrected and assembled into 9415 contigs using Canu ^34^ and consensus accuracy improved using Arrow from the PacBio software package. Total length of the Canu assembly is 1,848 Mb and N50 reached 816.8kbp, the longest contig was 12.8Mbp. We found that 6,937 contigs (~497Mbp) had relative read coverage less than 0.6, which may be from the heterogeneous diploid region of our fish sample, compared to 2,393 contigs (total length ~1347Mbp) with read coverage in the range of 0.6 to 1.8, most likely from the homologous regions (table 2) This is consistent with the 25-mer spectrum from our Illumina HiSeq2500 short read sequencing (figure 1). By summing all contig lengths normalized by read coverage, we determined the actual haploid genome size was at least 1.6Gbp. Contigs were aligned to self by using nucmer ^38^. 928 contigs contained in other contigs with low read coverage were removed, which was 27.3Mbp in total. All other contigs were retained.

#### Linkage Group Construction

RNA-seq data from two goldfish parents and their F_1_ offspring were download from NCBI (bioproject:PRJEB12518) ^21^. All reads were trimmed using Trimmomatic ^39^ (same configuring as in Gene Annotation) and aligned to the Canu assembly using hisat2 ^36^. Variant calling was performed via samtools mpileup and bcftools call (parameter ‘-m’) ^37^. We obtained ~5.6 M variants in total. SNPs with missing genotype or low read depth (<4) in more than 25% samples or with missing genotype in the two parents were removed (other filter: bcftools filter -g 10 -Ov -i ‘TYPE="snp" && QUAL>=10 && INFO/DP>=50’). SNPs that were homozygous in both parents or failing a Mendelian test were removed. We also required two SNPs on the same contig to be separated by at least 10Kbp. 14022 SNPs were kept after filtering and used for constructing genetic maps.

SNPs from the same contigs were grouped and ordered using ‘group’ and ‘seq.order’ from the R package ‘onemap’, with a LOD threshold of 5.5. Contigs with two or more groups (with each >= 3 markers) were broken at position with read depth valley and depth < 20 and depth < 20% quantile. In total, 16 contigs were broken. All SNPs were grouped using ‘group’ in the ‘onemap’ package. SNPs in each group were ordered using ‘seq.order’. Contigs were placed in each linkage group according to the ordered SNPs using chromonomer. After manual corrections, 50 long linkage groups were retained and named according their alignment to the zebrafish genome (LG1 and LG26 map to zebrafish chr1, LG2 and LG27 map to zebrafish chr2, and so on). Several short linkage groups, which were named according to their zebrafish alignment, were also retained. This assembly was named ‘carAur01’.

### Genome Annotation

#### Repeat Masking and Gene Structure Annotation

A custom repeat library for goldfish was built using RepeatModeler (http://www.repeatmasker.org/) based on the Canu assembly. Zebrafish and the custom repeat library were used to mask the genome by RepeatMasker (http://www.repeatmasker.org/, performed in MAKER3).

RNA-seq from seven goldfish tissues were performed to aid with gene annotation, include bone, brain (3 samples), eye, gill (2 samples), heart, muscle and tailfin. RNA libraries were prepared and sequenced on HiSeq2000 sequencer by NISC. All 2×125 pair-end reads were trimmed using Trimmomatic (ILLUMINACLIP:adapters/TruSeq3-PE-2.fa:2:30:10:8:true LEADING:3 SLIDINGWINDOW:20:20 MINLEN:40) and assembled via Trinity assembler without a genome-guide ^40^. All assemblies were clustered via CDHIT (-c 0.95 -aS 0.95 -uS 0.05), as EST evidence for Maker 3.0.

cDNA sequences from the Ensembl database (version 85, 69 species), NCBI vertebrate RefSeq and common carp (http://www.carpbase.org/gbrowse.php) were used as alternative RNA evidence. Proteins from the Ensembl database, common carp, and UniProt database (uniref90) were used as protein evidence. To annotate gene structure, we performed MAKER 3.0 ^41^ on the Canu assembly with Augustus prediction and the EST, RNA, protein evidence. Gene structures were lifted over to the carAur01 assembly using liftover ^42,43^ or crossmap (https://sourceforge.net/projects/crossmap/files/).

Because our fish was not fully homozygous, we needed to identify those genes in the heterozygous diploid regions. All cDNA sequences from Maker gene models were aligned to self by megablast. Alignments with identity ≥ 97.5% and coverage of both sequences ≥ 70% were kept. Alignments were retained if they satisfied one of the following restrictions: (1) identity >= 99.5% and the relative coverages of both contigs where the two genes were located were less than 0.8, (2) the relative coverage of both contigs was less than 0.75, (3) the relative read coverage of either contig was less than 0.6. DNA sequences from all remaining aligned genes were fetched and aligned using lastz and chained with axtChain. All alignments with matched basepairs covering less than 0.6 of both genes or with identity less than 95% were discarded. Only the shorter of the two genes in the retained alignments was masked and not used for following analysis.

MAKER3 generated 81,778 coding gene models, of which 80,062 were liftover’ed to carAur01, and 9,738 genes were masked as one allele of the heterozygous genes. The average exon and intron length was ~202bp and ~174bp. The distribution of exon and intron size is similar to zebrafish, grass carp and common carp (supplemental figure 2).

#### Non-coding RNA annotation

Non-coding RNA sequences from other species were downloaded from NONCODE ^44^ (zebrafish and human), RNAcentral ^45^ and Ensembl ncrna (ver. 85) ^46^. All sequences were first aligned to the genome using blastn in the NCBI-BLAST+ package ^47^ (-evalue 1e-4 – perc_identity 80). All genomic target regions were fetched and refined using exonerate^48^ for each query. Exonerate alignments for each query RNA were kept if they satisfied: (1) score ≥ 0.9 best score for the query; (2) query coverage ≥ 0.6; (3) query identity ≥ 0.7; (4) non-canonical splice site ≤ 3.

Trinity genome-guided assembly was performed on the RNA-seq data from the seven tissues. ‘align_and_estimate_abundance.pl’ from the Trinity package was used to estimate the expression of each transcript. Transcripts with expression lower then 1 TPM were filtered. All remaining transcripts were aligned to the Canu assembly using the same BLASTN-exonerate approach except using a higher identity 90%. Exonerate alignments for each query RNA were kept if they satisfied: (1) score ≥ 0.95 best score for the query; (2) query coverage ≥ 0.75; (3) query identity ≥ 0.9; (4) non-canonical splice sites ≤ 3. All Trinity transcripts with no alignment to any MAKER genes or with Trinotate PFam/Spot annotation were also removed ^49^. Coding potential of the remaining transcripts were predicted by using CPC ^50^. Transcripts with ‘coding’ labels were removed. All the remaining exonerate results were transformed to GFF3 and merged using ‘cuffcompare’ from cufflinks package.

Hairpin sequences from miRBase were also aligned to the genome using the BLASTN-exonerate approach. Alignments were retained if they satisfied: (1) score ≥ 0.9 best score for the query and (2) query coverage >90%, identity >90%.

The genome was scanned against the Rfam database using cmscan from the Infernal package (version 1.1.1) ^51,52^. Only hits with bit score ≥ 30 and E-value ≤ 10e-6 were kept. When dealing with overlapping hits, we kept the hit amongst all overlapping hits that had the highest bit score.

#### Conserved Noncoding Elements (CNE) Identification

All-to-all pairwise genomic alignment was performed using lastz (--gapped -- ambiguous=n --step=3 --strand=both --masking=100 --maxwordcount=100 -- identity=70..100 --format=axt) and axtToChain for four species (goldfish, common carp, grass carp, zebrafish) and transformed to pairwise MAF format and split at gaps longer than 30bp (chainToAxt –maxGap=30, then axtToMaf -score). All the pairwise MAF files were transformed to multiple alignment MAF files using roast (P=multic). Phylogenetic models were fit for each chromosome, linkage group or scaffold using phyloFit (--tree ‘(ZF,(GC,(GF,CC)))’ --subst-mod REV --nrate 4), which was used by phastCons for computing conservation scores and most conserved regions. The most conserved regions out of exons (of coding or noncoding genes) were defined as CNE (conserved noncoding element). goldfish (or common carp) CNE that overlapped the goldfish-goldfish (or common carp-common carp) self chain-net alignment regions were retained either as both WGD copies or as singletons.

#### Gene Functional Annotation

Interproscan5 ^53^ was used to annotate the Interpro/GO/Pathway function for all protein-coding genes.

#### SNV and DIV

2×250 read pairs from a second gynogenic goldfish (GF71, 73X coverage) and a wild-type goldfish (WTGF, 70X coverage) were aligned to the carAur01 assembly using bwa mem (bwa mem -t 16 -I 538.,149.3). Most Probable Genotype (MPG) (https://research.nhgri.nih.gov/software/bam2mpg/index.shtml, https://github.com/nhansen/bam2mpg) ^54^ was used to call variants from the bwa mem produced bam files. The MPG output variant calls were converted to VCF for variants with a minimum Most Probable Variant (MPV) score of 10 or greater with a MPV-score/read-coverage ≥0.5

#### Functional Enrichment

Fisher exact tests were performed to identify significantly enriched GO molecular functions among goldfish, common carp, grass carp and zebrafish. We also performed the same tests between duplicated retained genes and single-copy-lost genes in goldfish for each GO terms in the ‘molecular function’ and ‘biological process’ domain (figure 9). Compared to the other three species, goldfish show enriched function in channel activity and depressed function in olfactory receptor activity (figure 10).

### Evolution Analysis

#### Ohnolog Gene Clusters

Protein and cDNA sequences of zebrafish (GRCz10) were downloaded from the Ensembl database. Grass carp sequences were downloaded from Grass Carp Genome Database (GCGD) ^55^. Common carp sequences were downloaded from NCBI (GCF_000951615.1).

We performed all-to-all Blastn on the cDNAs from the four species. Non-overlapping alignments from the same cDNA pairs were concatenated. We identified synteny blocks for each pair of species through iteratively merging nearby aligned gene pairs with, at most, five unaligned genes between them. Alignments were used as an edge to group genes into clusters with constrained gene numbers for each species according to whether it was before or after the carp WGD event (zebrafish:grass carp:common carp:goldfish = 1:1:2:2). Two genes or gene clusters were merged if the number of edge between them was > 50%N_1_N_2_, or > 20%N_1_N_2_ and there were edges linked between the two genes to a matching outgroup gene according to the species tree ‘(zebrafish, (grass carp, (common carp, goldfish)))’, where N_1_ and N_2_ were the number of genes in each gene cluster. The priority for the edge for aggregate genes or gene clusters were edges in synteny blocks and then ‘reciprocal best hit’ edge. Other edges were used to rescue and merge some genes into those non-full-size (i.e. 1:1:2:2) clusters.

#### Phylogenetic Analysis

Proteins from all 1:1:2:2 ohnolog clusters were multiple aligned using MAFFT ^56^ with ‘-- auto’ option, then transformed to codon alignment using ‘tranalign’ from EMBOSS Suite ^57^. Poorly aligned codon regions were eliminated using Gblocks ^58^. The third position of all codons was filtered out into separated alignments. All third-codon sequences from the same chromosomes were concatenated for building phylogenetic trees. ML tree was built using RAxML ^59^ with the model GTRGAMMA. Pairwise synonymous substitutions were computed by using ‘codeml’ from the PALM package (runmode = −2, method = 0) ^60^. Divergence time of the carp WGD event was estimated by 20.5*L(WGD)*2/L(grass_carp,carp), where 20.5 is the divergence time of grass carp and common carp in unit Mya, L(WGD) is the average branch length from WGD event to goldfish and common carp, L(grass_carp, carp) is the average branch length between grass_carp and common carp or goldfish. Similar estimation was performed for the speciation of common carp and goldfish.

#### Expression Comparison between Retained WGD Gene Pairs

Co-linear blocks were fetched from the goldfish self chain-net alignment. Gap larger than 20kbp was broken. Blocks shorter than 50kbp were removed. Blocks were removed if it overlaps other longer blocks. The two sequences in each collinear block were presumed to be derived from the same sequence before the carp WGD event. WGD gene pairs were fetched from these collinear blocks for follow-up analysis. Exons or CNEs that were lost in exactly one sequence from each block were also identified. The genes that CNE were predicting to regulate were defined as the nearest gene(s) in 5kbp windows on both sides.

RNA-seq reads from the seven tissues were mapped to the carAur01 assembly using STAR (default settting and two pass). Expression levels (TPM) were estimated using RSEM (rsem-calculate-expression --paired-end --forward-prob 0.0 --alignments -p 16 -- seed 987347 --calc-ci --calc-pme --estimate-rspd --time --no-bam-output) and transformed to logTPM=log2(TPM+1). Euclidean distances or correlation coefficients of the expression between WGD gene pair were calculated in R. 449 gene pairs were silenced, another 649 gene pairs contained exactly one silenced gene. The remaining 19,500 genes (9,750 gene pairs with both genes expressed) were hierarchically clustered using the ‘hcluster’ and ‘ward.D2’ method in R, based on the logTPM value and Euclidean distance. Tissue specific expressed gene pair was defined as gene pair with TPM>=4 in one gene and TPM<0.5 in the other gene in at least one tissue. Expression standard deviations across the seven tissues were also calculated for each gene.

Gene pairs were divided into 6 groups according to their pairwise cDNA identity (≤86%, 86-88%,88-90%,90-92%,92-94%,>94%). Histogram of expression distances for each group were computed in R using ‘hist’ with bin size 2. In order to illuminate the relationship between exon loss and expression distance, gene pairs were divided into 4 groups: no exon loss, one exon loss, two exon losses, three or more exon losses. One sided Wilcoxon rank sum tests were performed for each pair of groups. For CNE lost, Wilcoxon rank sum test was performed on the expression standard deviation between genes in the no-CNE-lost group and those in the CNE-lost group, using only gene pairs with CNE loss but no exon loss.

In order to find out which biological functions were prone to diverging after the WGD, we performed Wilcoxon rank sum tests on the expression distance between genes inside the GO terms and genes outside the GO terms. The top 20 and bottom 20 GO terms with p < 0.1 were plotted in figure 15.

### Software and Databases

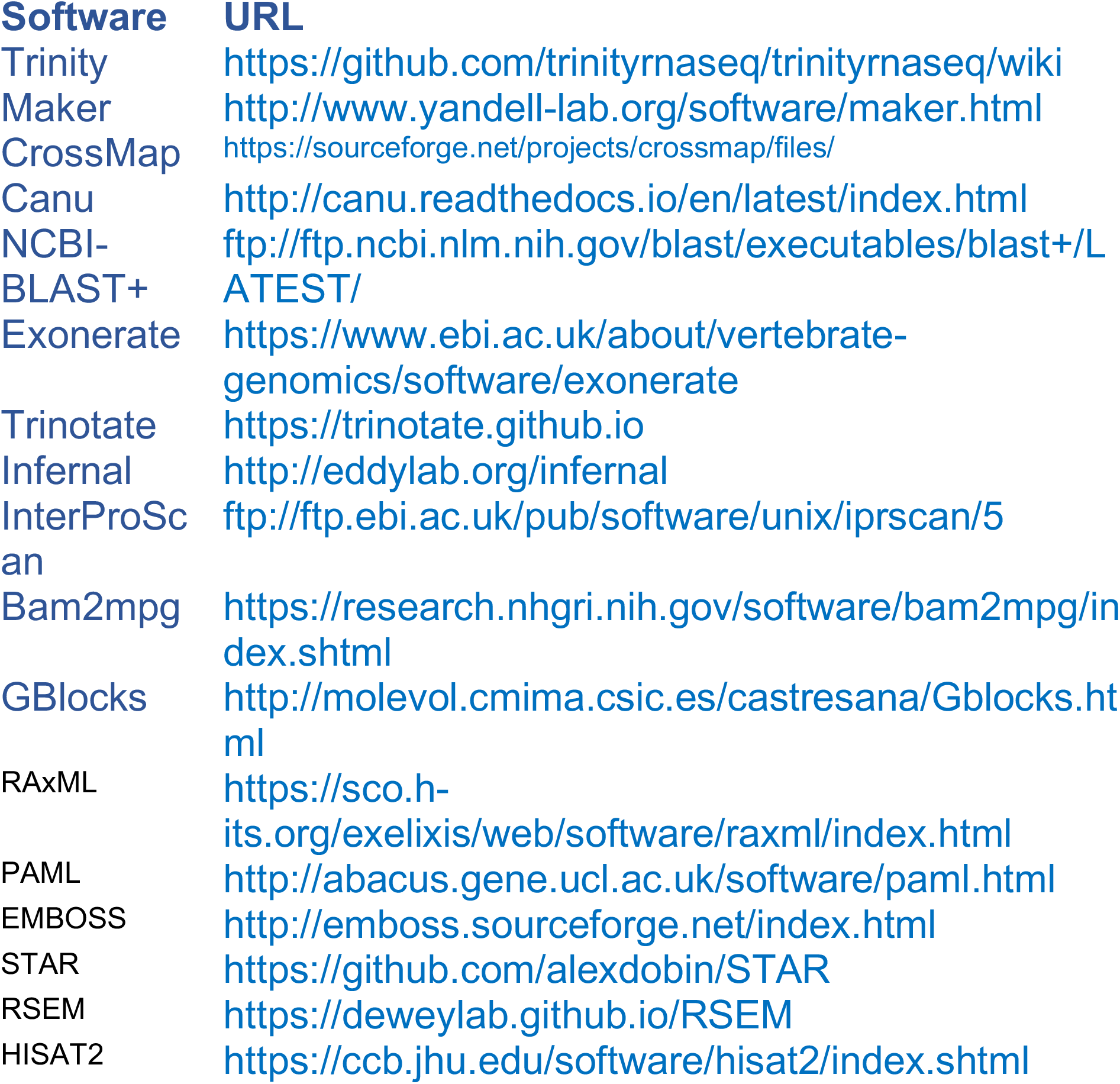

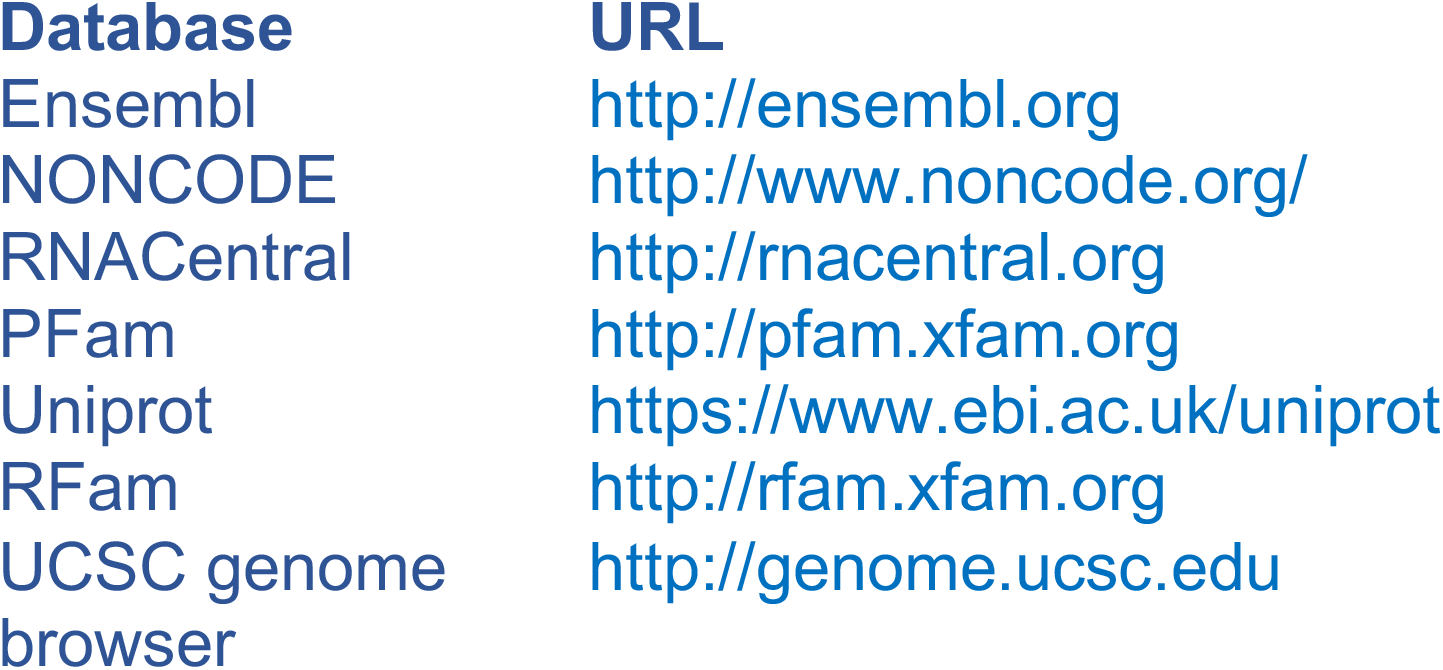

## Supplemental data

### Supplemental Tables

**Supplemental Table 1.** Pacbio read statistics

|  | Raw Reads | Corrected Reads |
| --- | --- | --- |
| Counts | 16,671,136 | 11,884,085 |
| Mean length (bp) | 7,800 | 6,810 |
| Coverage | ~71 | ~45 |
| Peak length (kbp) | ~9.8 | ~8.0 |

**Supplemental Table 2.**
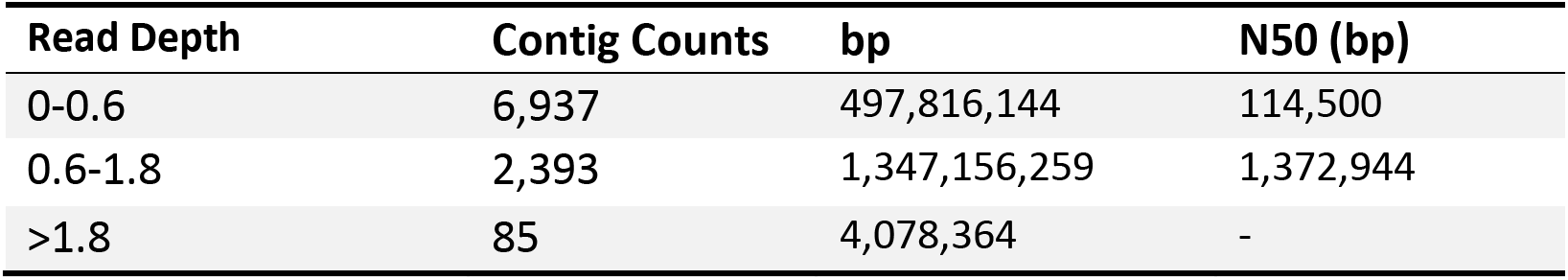
Assembly statistics for different coverage groups

| Read Depth | Contig Counts | bp | N50 (bp) |
| --- | --- | --- | --- |
| 0-0.6 | 6,937 | 497,816,144 | 114,500 |
| 0.6-1.8 | 2,393 | 1,347,156,259 | 1,372,944 |
| >1.8 | 85 | 4,078,364 | - |

**Supplemental Table 3.** Repeated DNA statistics

|  | **Goldfish** | **Common Carp**<sup>23</sup> | **Zebrafish**<sup>27</sup> |
| --- | --- | --- | --- |
| **Total base pairs** | 721,087,053<br>(39.6%) | 672,246,354<br>(39.2%) | 715,370,858<br>(52.24%) |
| **DNA transposon** | 16.38% | 17.53% | 34.3% |
| **LTR** | 4.89% | 4.35% | 5.07% |
| **LINE** | 4.50% | 4.90% | 2.83% |
| **SINE** | 0.47% | 0.47% | 2.34% |
| **Satellite** | 1.27% | - | 1.78% |
| **RC** | 1.89% | - | 0.94% |
| **Simple** | 3.27% | - | 4.12% |
| **Unknown** | 6.88% | - | 0.34% |

The breakdown of the various repeat elements presented in goldfish, common carp, and zebrafish. The percentage of the total genome is indicated in parentheses. The larger fraction of DNA transposons in zebrafish is responsible for its significantly larger size compared to the pre-duplication carp or goldfish genomes.

**Supplemental Table 4.** Core eukaryotic genes using Benchmarking Universal Single-Copy Orthologs (BUSCO)

|  | **Goldfish** | **Common carp** | **Zebrafish** |
| --- | --- | --- | --- |
| **Complete BUSCOs** | 4,204 | 3,828 | 4,384 |
| **Complete and single-copy BUSCOs** | 1,990 | 1,695 | 4,145 |
| **Complete and duplicated BUSCOs** | 2,214 | 2,133 | 239 |
| **Fragmented BUSCOs** | 257 | 436 | 113 |
| **Missing BUSCOs** | 123 | 320 | 87 |
| **Total BUSCO groups searched** | 4,584 | 4,584 | 4,584 |

Using the “Benchmarking of Universal Single-Copy Orthologs” *Actinopterygii* gene set, we determined the goldfish genome assembly has 97.3% of the BUSCO in at least one copy (91.7% complete BUSCO genes, 5.6% fragmented, and 2.7% missing) with 48.3% complete in both copies, compared to the common carp assembly which has 83.5% complete BUSCO, 9.5% fragmented, 7% missing and 46.5% complete with both gene pairs represented.

### Supplemental Figures

**Supplemental Figure 1.**
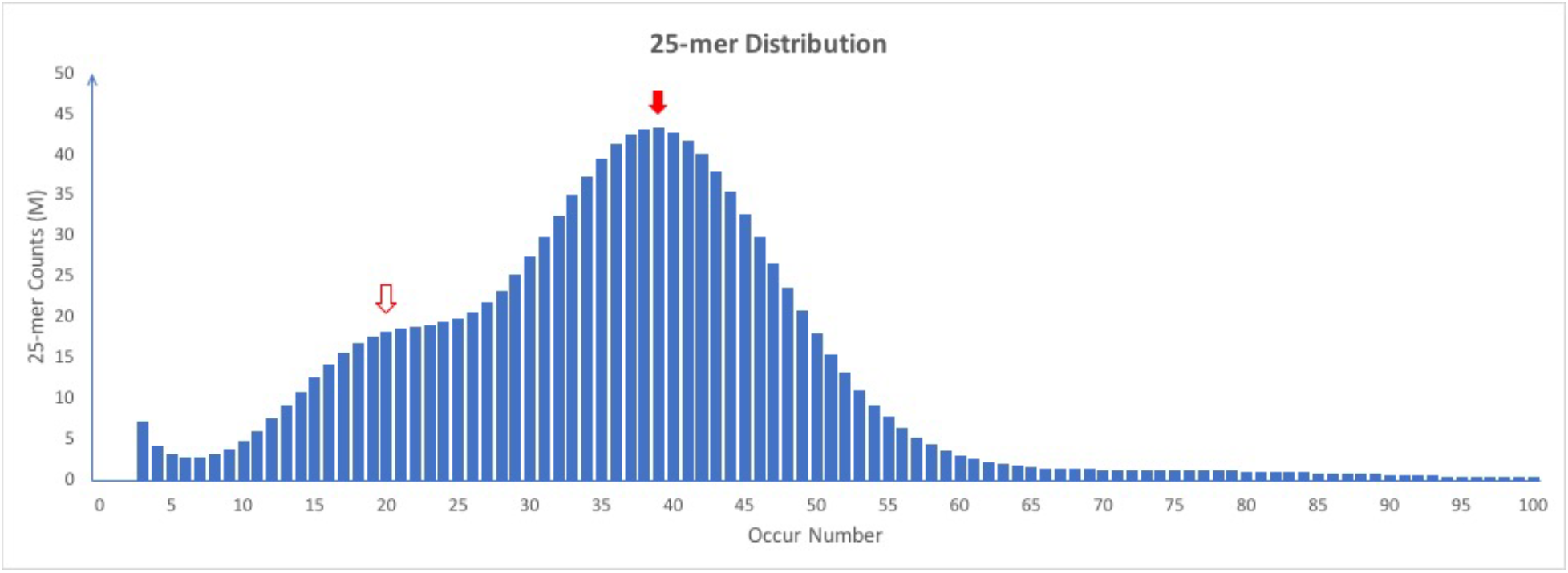
25-mer occurrence distribution from 2 × 125 bp Illumina paired-end reads. The two peaks indicate that a fraction of the genome was not sequenced to the same depth of coverage, i.e. part of the genome (approximately 16% from The Canu assembly) was at 20X coverage instead of 40X (white arrow *vs*. red arrow). The 20X peak was indicative of regions of the genome that were not homozygous.

**Supplemental Figure 2.**
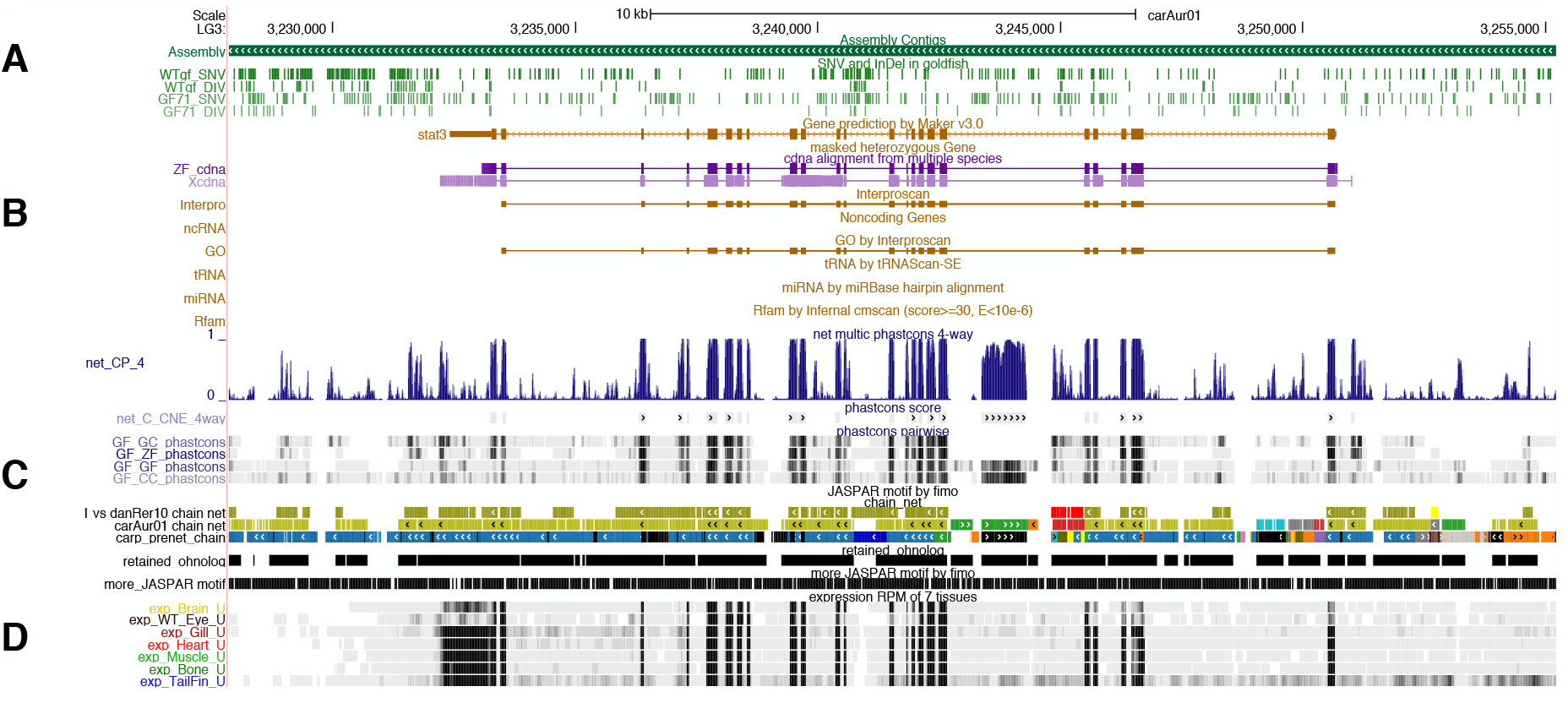
Screenshot of the UCSC Genome Browser implementation of the carAur01 assembly. Genome annotation includes: A) Assembly, SNV and DIV data from sequencing three “wild-type” Wakin goldfish, B) gene model annotation C) multiple genome alignment tracks that compare goldfish to zebrafish, grass carp, and common carp to identify conserved coding and non-coding (i.e. enhancers/promoters) sequences, D) gene expression from 7 adult goldfish tissues. Hub available at: https://research.nhgri.nih.gov/goldfish/

**Supplemental Figure 3.**
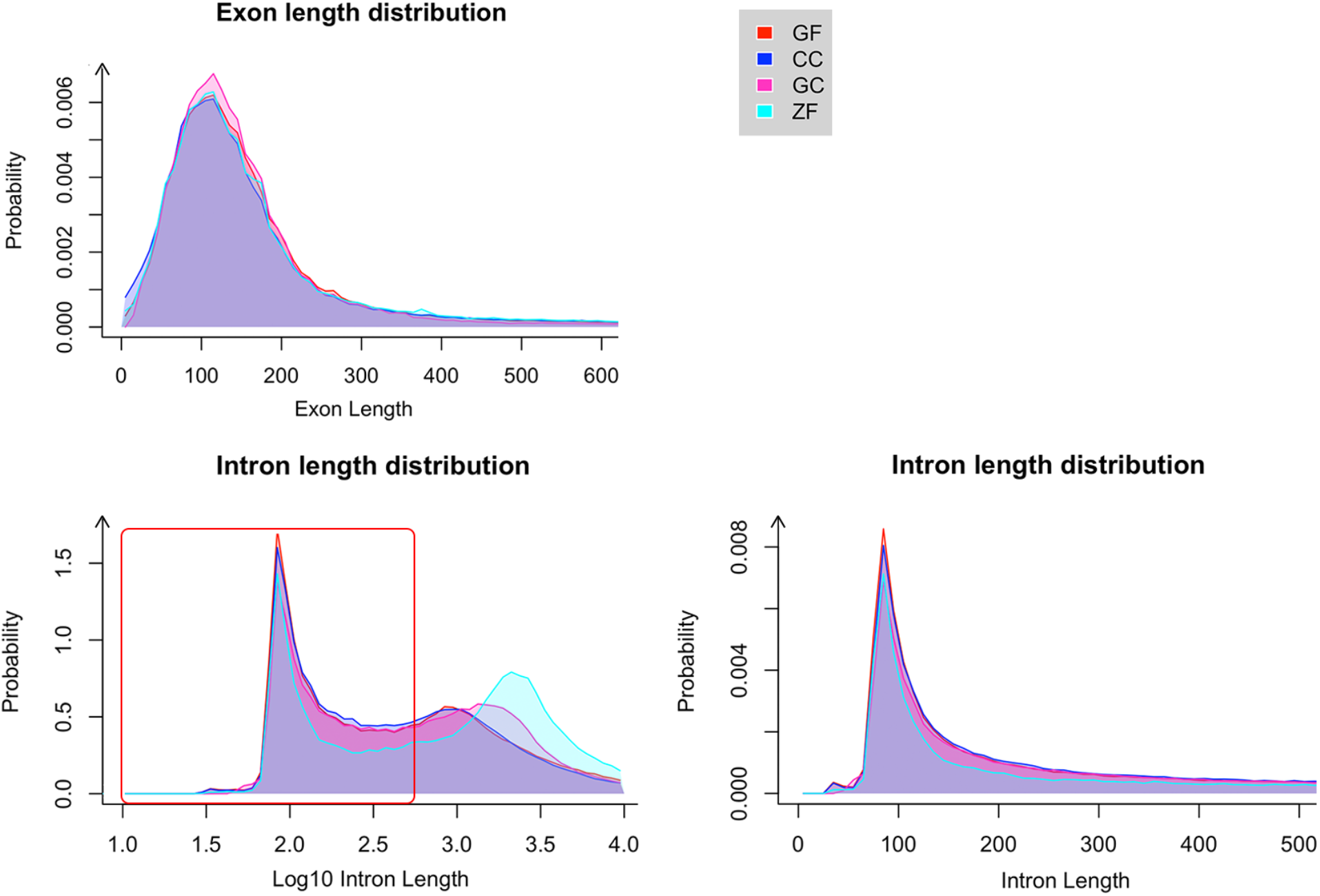
Distribution of exon and intron lengths. Bottom right panel is an enlargement of the red box in the bottom left panel. GF: goldfish, CC: common carp, GC: grass carp, ZF: zebrafish.

**Supplemental Figure 4.**
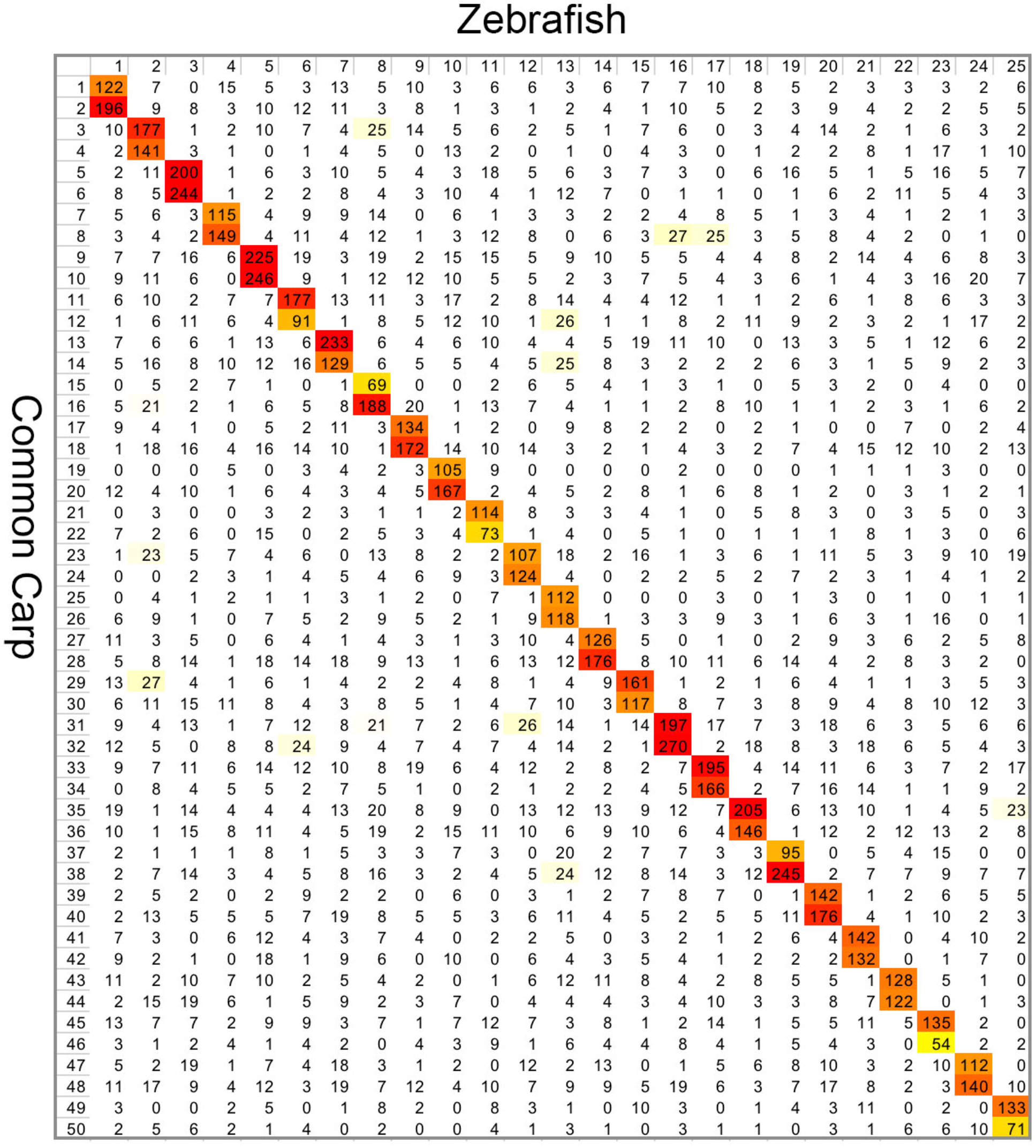
Reciprocal best hit (RBH) gene counts between zebrafish and common carp chromosomes. Red to yellow indicates high to low numbers.

**Supplemental Figure 5.**
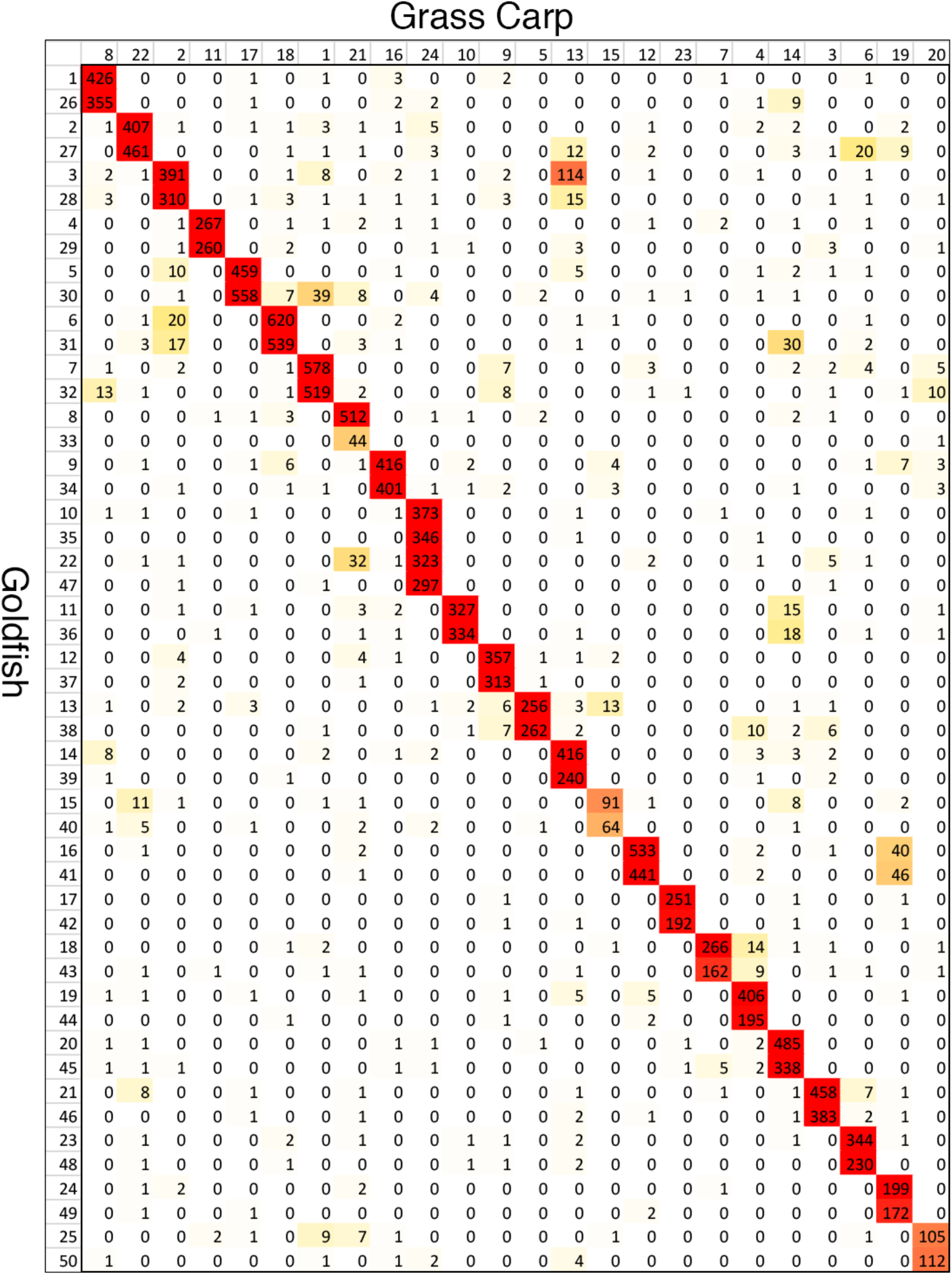
RBH gene counts between grass carp and goldfish chromosomes. Red to yellow indicates high to low numbers.

**Supplemental Figure 6.**
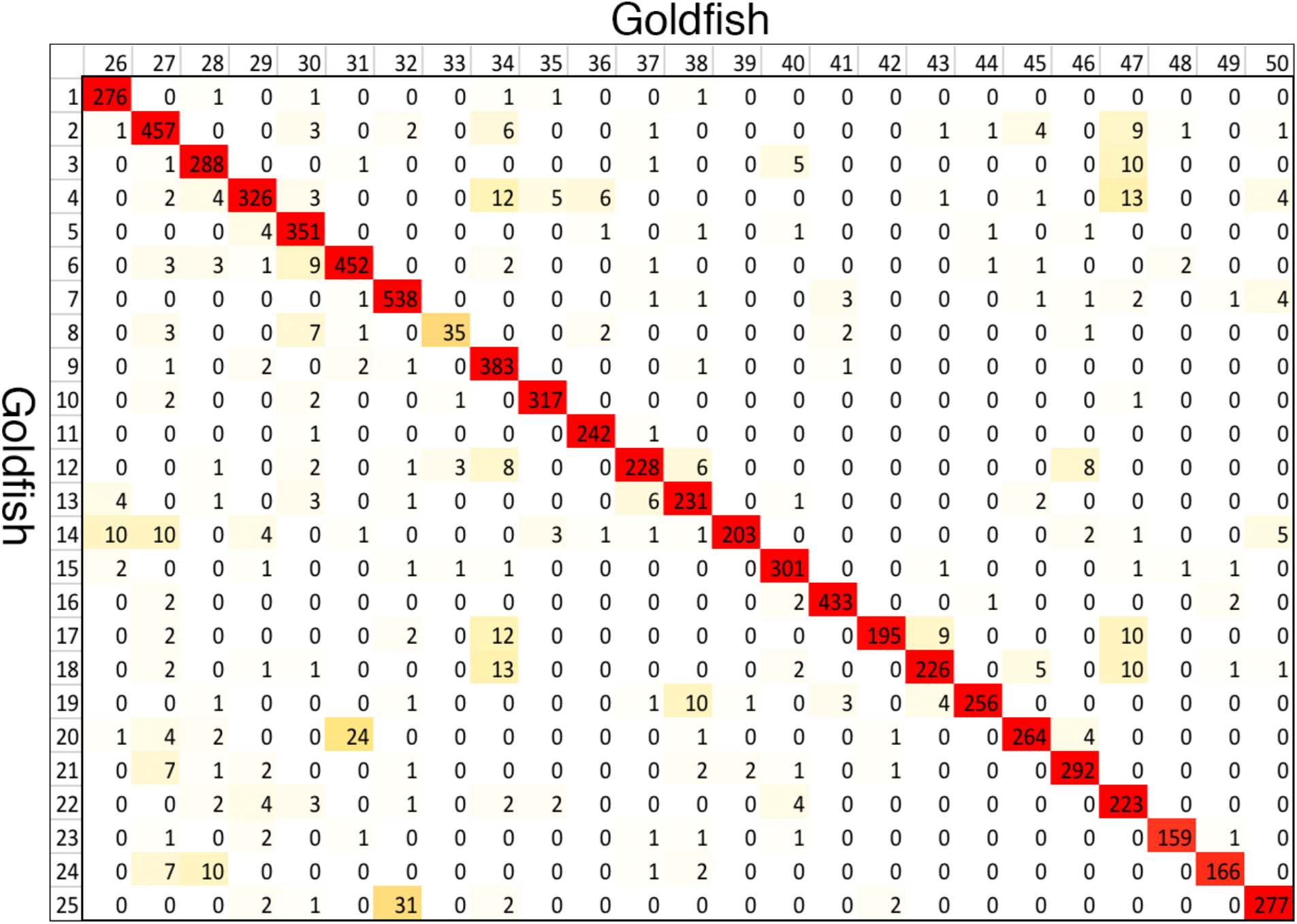
RBH gene counts between goldfish whole genome duplicated chromosomes. Each row or column is one chromosome. Red to yellow indicates high to low numbers.

**Supplemental Figure 7.**
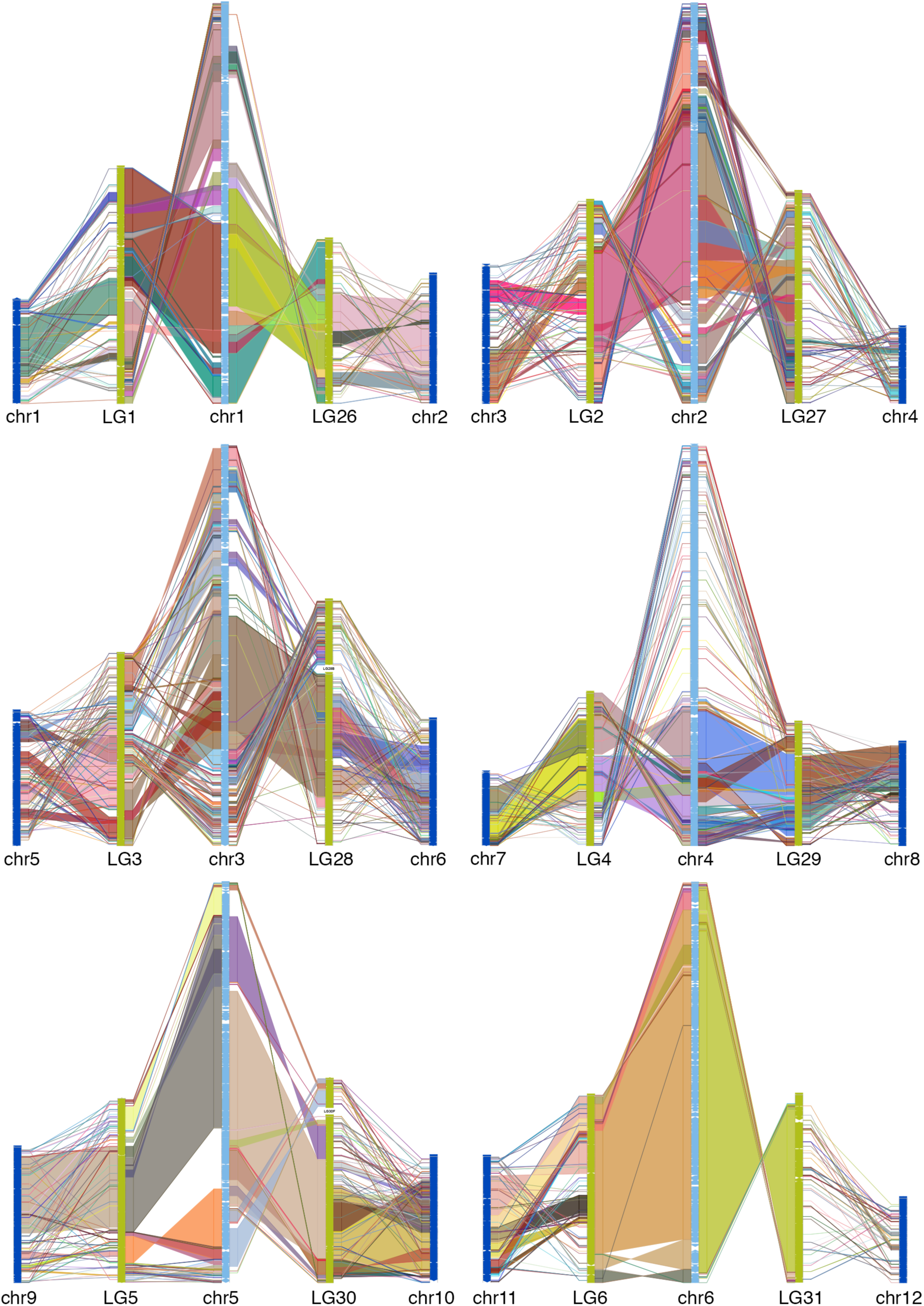
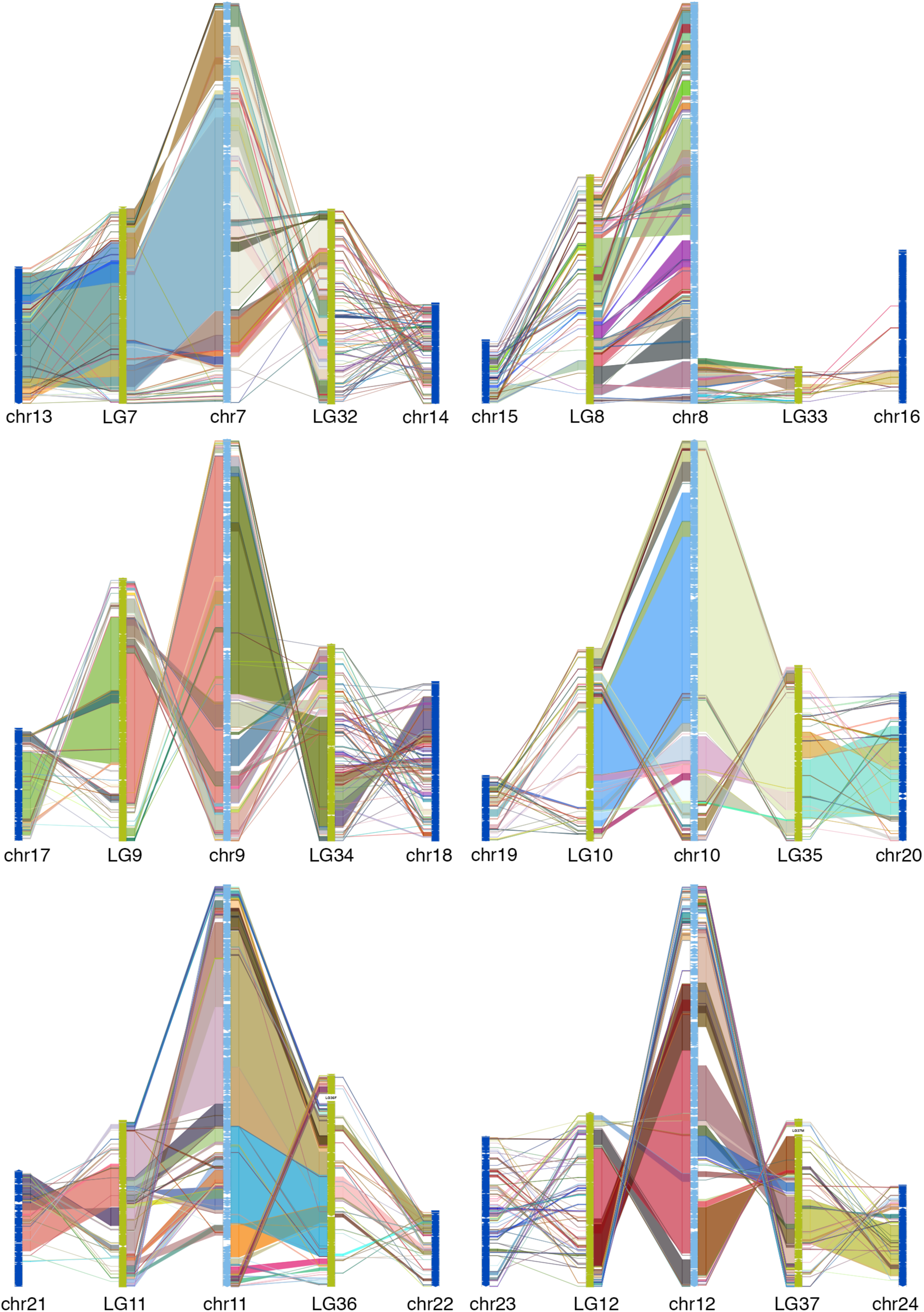
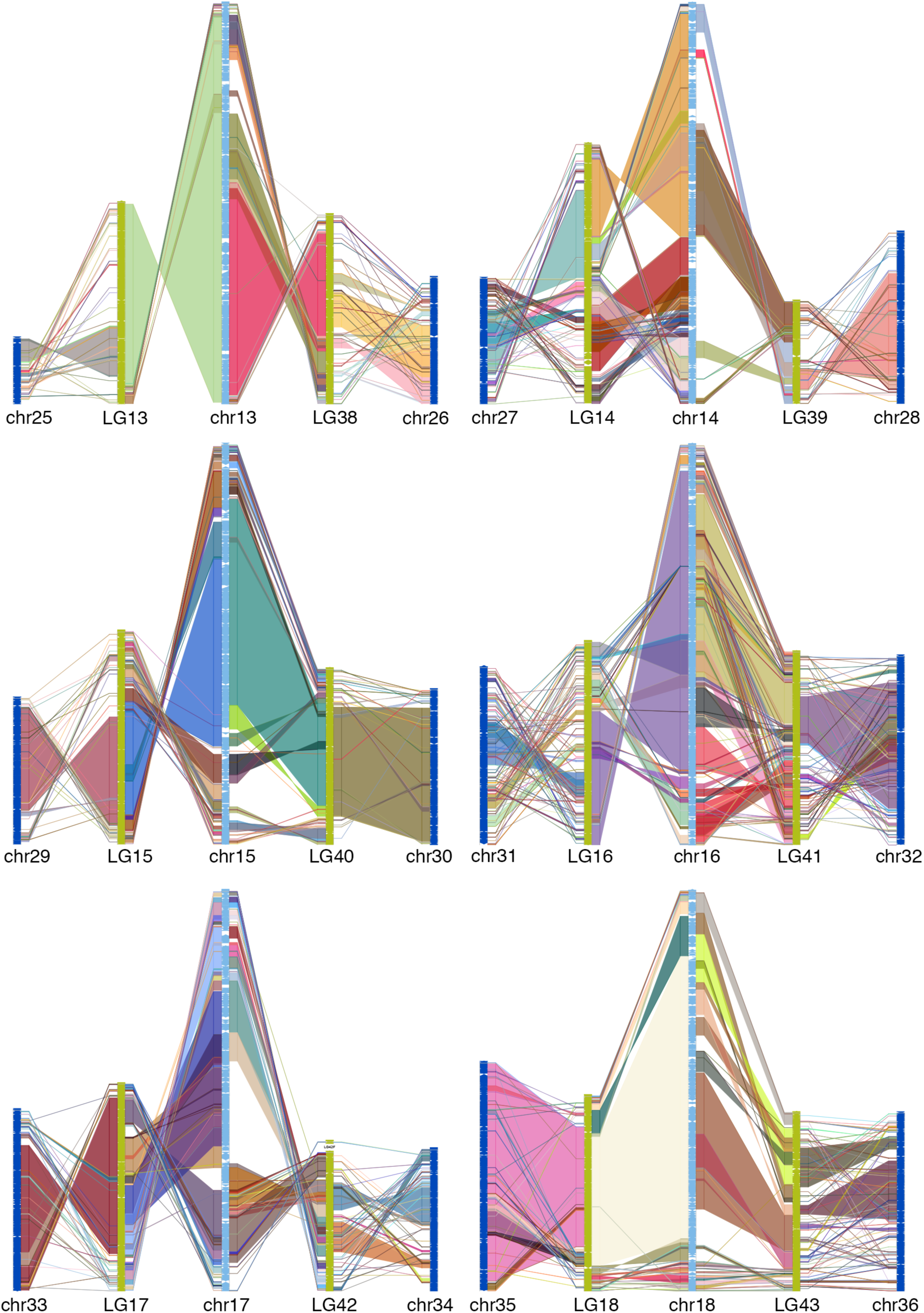
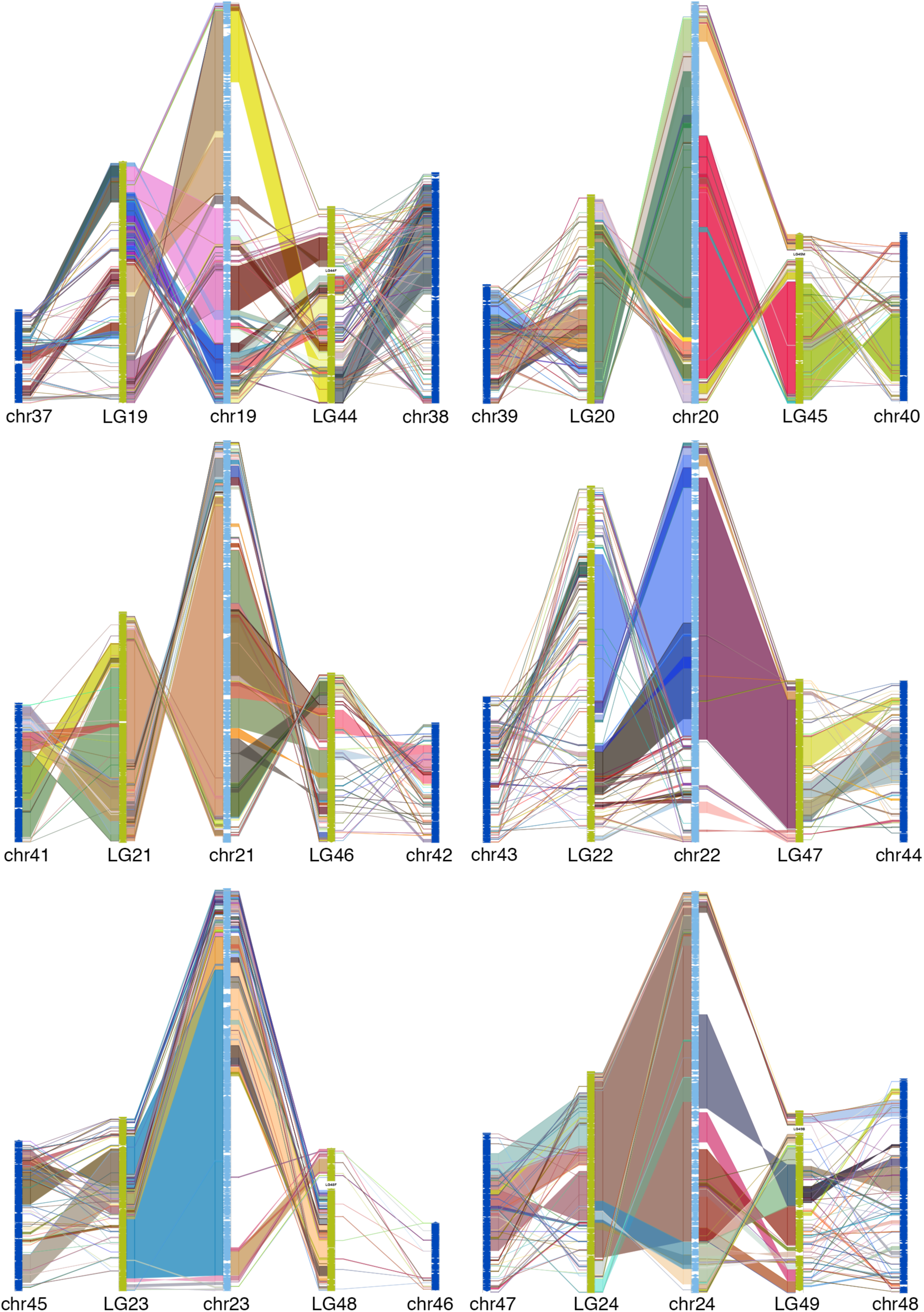
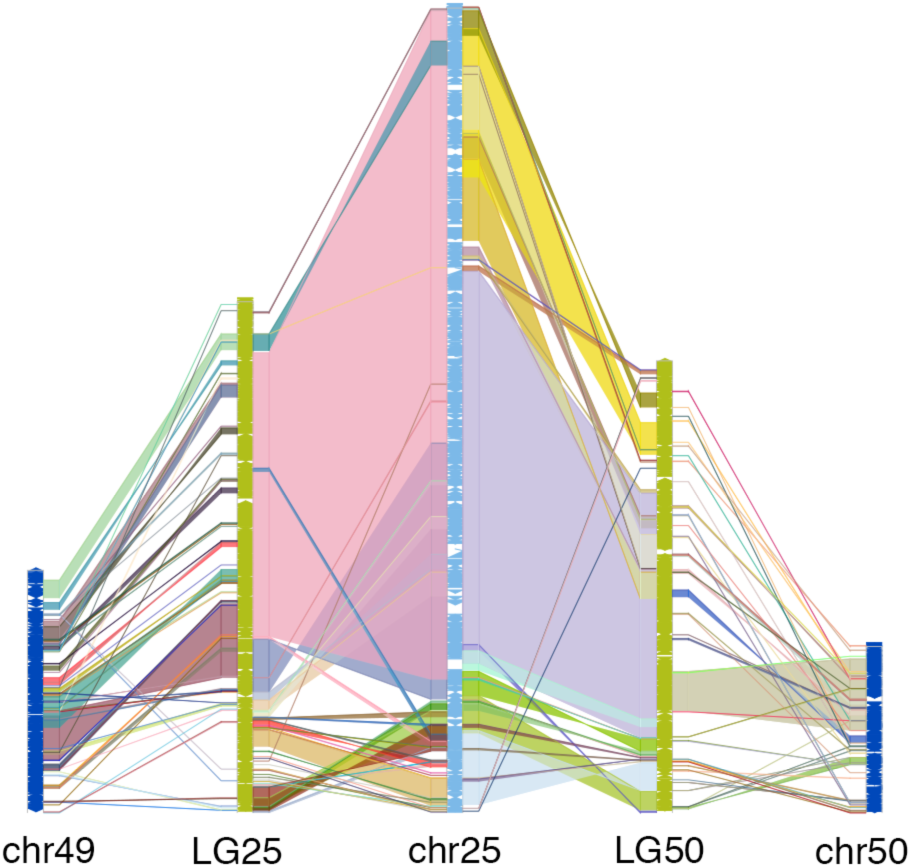
Chain-Net alignment between each zebrafish chromosome (middle light blue bars) and two corresponding whole genome duplicated goldfish chromosomes (green bars), and goldish to common carp (blue bars). Lines or blocks between bars show alignments between the two chromosomes. Typically one of goldfish chromosome pairs contained a significantly larger block of conserved col linearity than the other, but both chromosomes show remarkable stability across 60 million years of evolution.

**Supplemental Figure 3.**
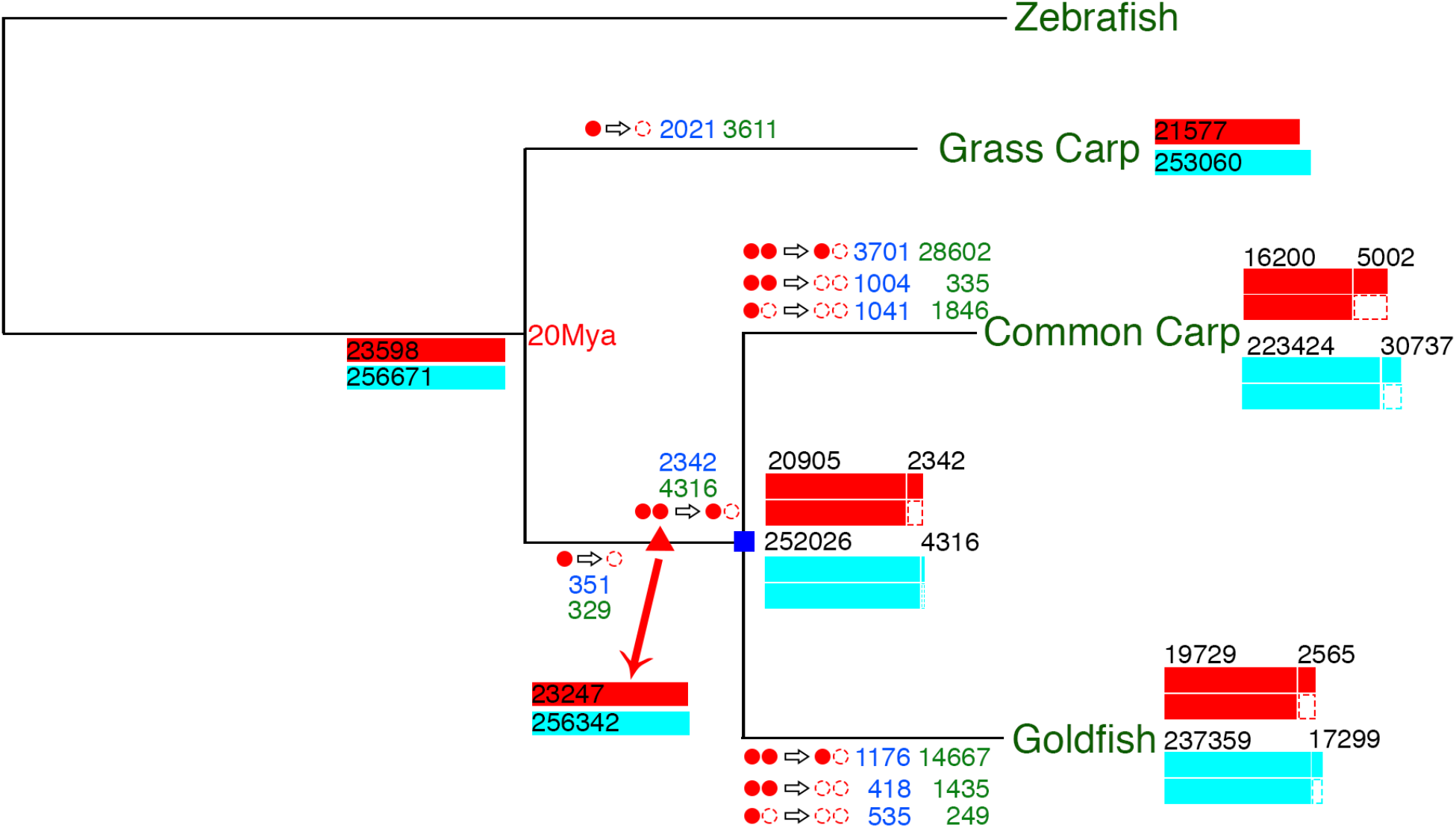
Gene and CNE lost in phylogenetic history. Using zebrafish as the reference, the tree tracks gene and CNE loss at different evolutionary branchpoints. On each branch, a filled red circle indicates a retained copy, an open red circle indicates a lost copy. Numbers in blue are for lost genes, numbers in green are for lost CNEs. Red (light blue) boxes: retained genes (CNEs), scaled by percentage. The black number over each box is the number of retained genes or CNEs. The red triangle represents the carp whole genome duplication event at 14.4 Mya. The blue square marks the speciation of common carp and goldfish at 11.0 Mya. Maximum likelihood phylogenetic tree was constructed by using the third position of all codons of ohnolog genes. It is clear to see that rates of gene and CNE loss accelerated after the genome duplication event. We assume most cases where both copies of a gene were lost in either goldfish or carp, this loss occurred after separation from grass carp but before the whole genome duplication.

**Supplemental Figure 4.**
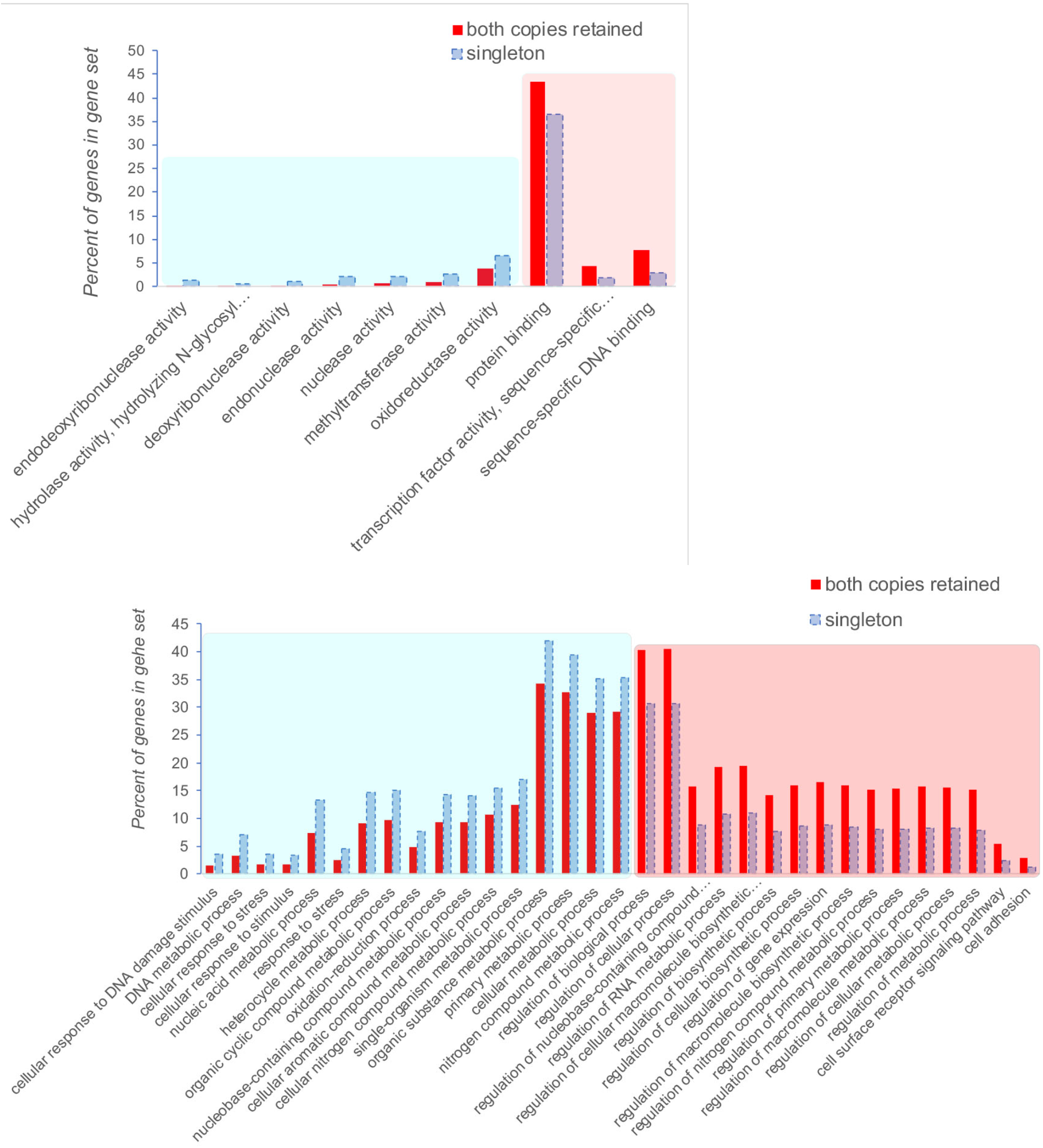
GO terms prone to retaining both gene copies (blue rectangle) or losing one copy (blue rectangle) after whole genome duplication in goldfish. Zebrafish was used as the reference genome (FDR<0.01). Upper: GO molecular functions. Lower: GO biological processes. “Percent of genes in gene set” describes how many genes in each class (both preserved or one copy lost) fall into each GO term, i.e. are some genes in each class over-represented (more likely or less likely to be lost) compared to neutral.

**Supplemental Figure 5.**
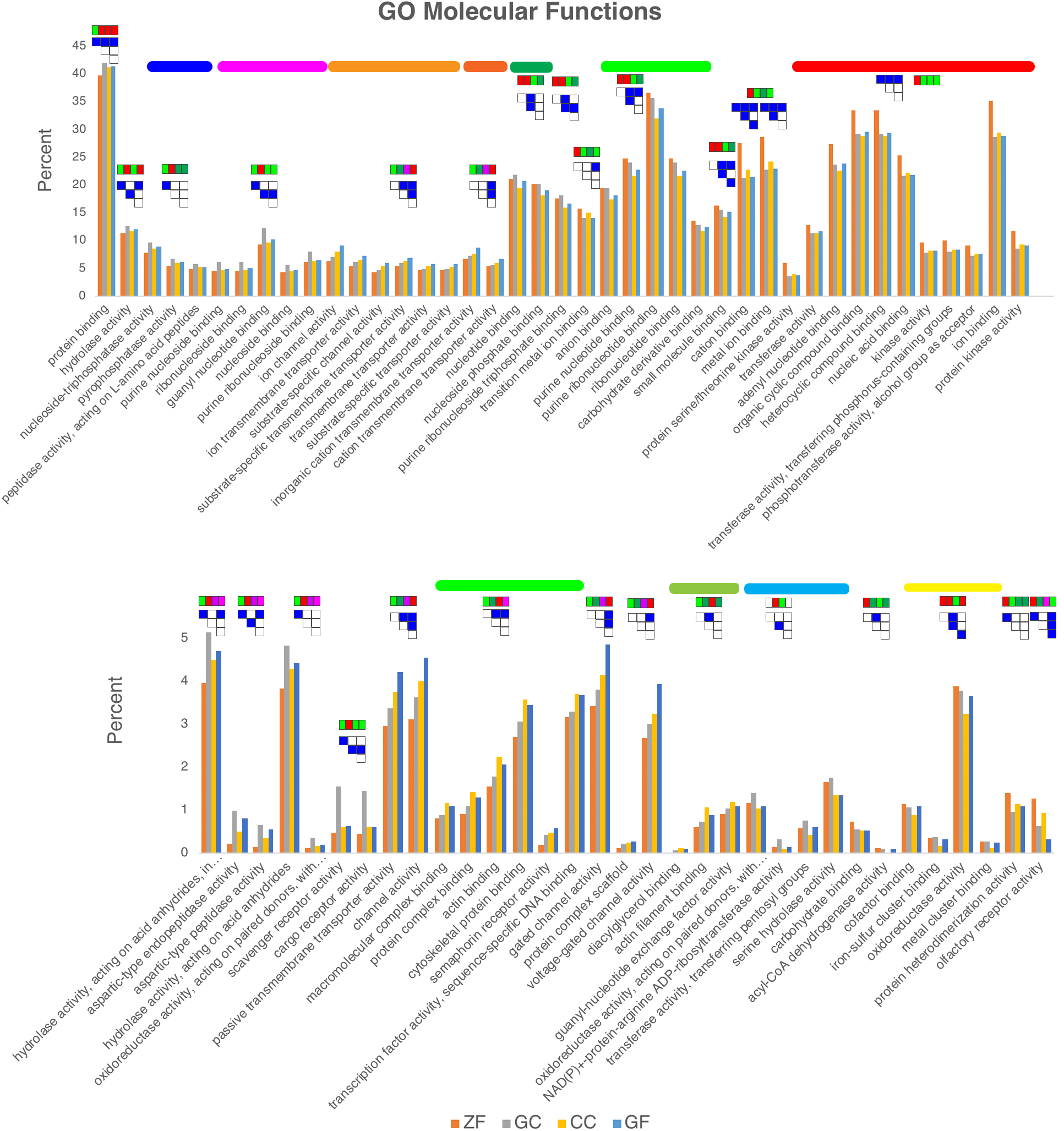
GO molecular function comparison among zebrafish(ZF), grass carp (GC), common carp (CC), goldfish (GF). The histogram shows the percentage of genes in the gene set. The four colored boxes indicate the relative values among the four species, green for low, red for high, pink, purple or dark green show middle values from higher to lower. The blue or white matrix indicates pair-wise significant values, blue for significant (p-value<0.01 and FDR<0.1), white for non-significant. Color bars indicate clusters with similar trends among the four species.

**Supplemental Figure 11. a.**
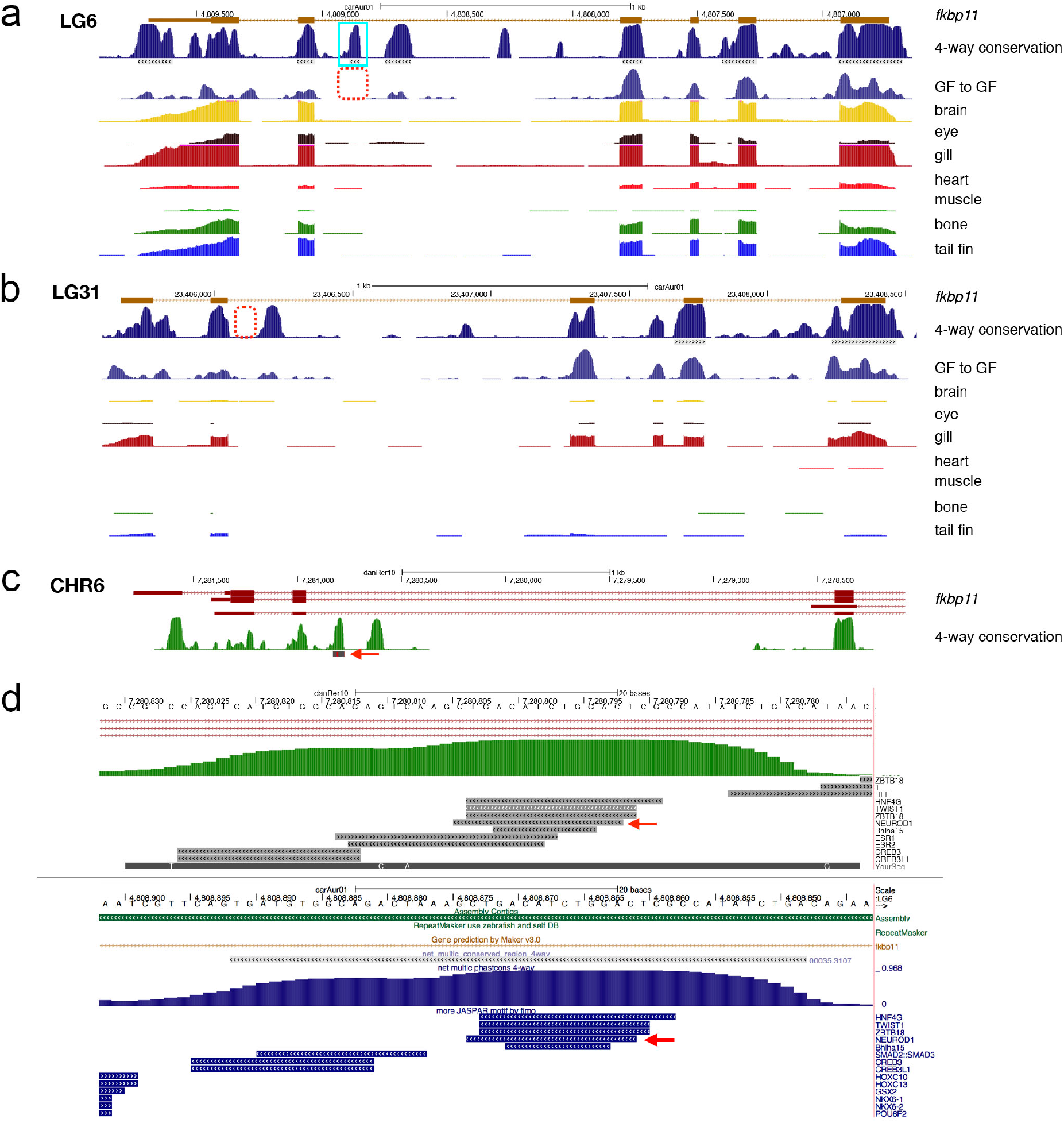
Screenshot example of the *fkbp11* gene containing conserved, non-coding elements on linkage group 6. The “4-way conservation” peaks are from comparing goldfish, zebrafish, common carp and grass carp, gray bars beneath the peaks are regions satisfying the criteria for CNE. The GF to GF track shows sequences conserved in both chromosomal duplicates. The red dotted box shows the missing sequences on the matching duplicated chromosome (LG31). The remaining tracks are the RNA-seq data from each tissue, showing strong expression in brain, eye, gill, bone, and tail fin, with weaker expression in the muscle and heart. **b.** The region on LG31 containing the second copy of *fkbp11*. The red box shows where the missing CNE should be. Expression levels for most of the tissues is very low with the exception of expression in the gill. **c.** Zebrafish fkbp11 showing the 4-way conservation peaks and the BLAT hit using the goldfish sequences from LG6 (red arrow). **d.** Magnified view of the zebrafish CNE (upper) and goldfish CNE (lower) including JASPAR-predicted transcription factor binding sites. Red arrow marks a highly conserved *neurod1* site, a potentially strong enhancer for brain and eye expression.

**Supplemental Figure 6.**
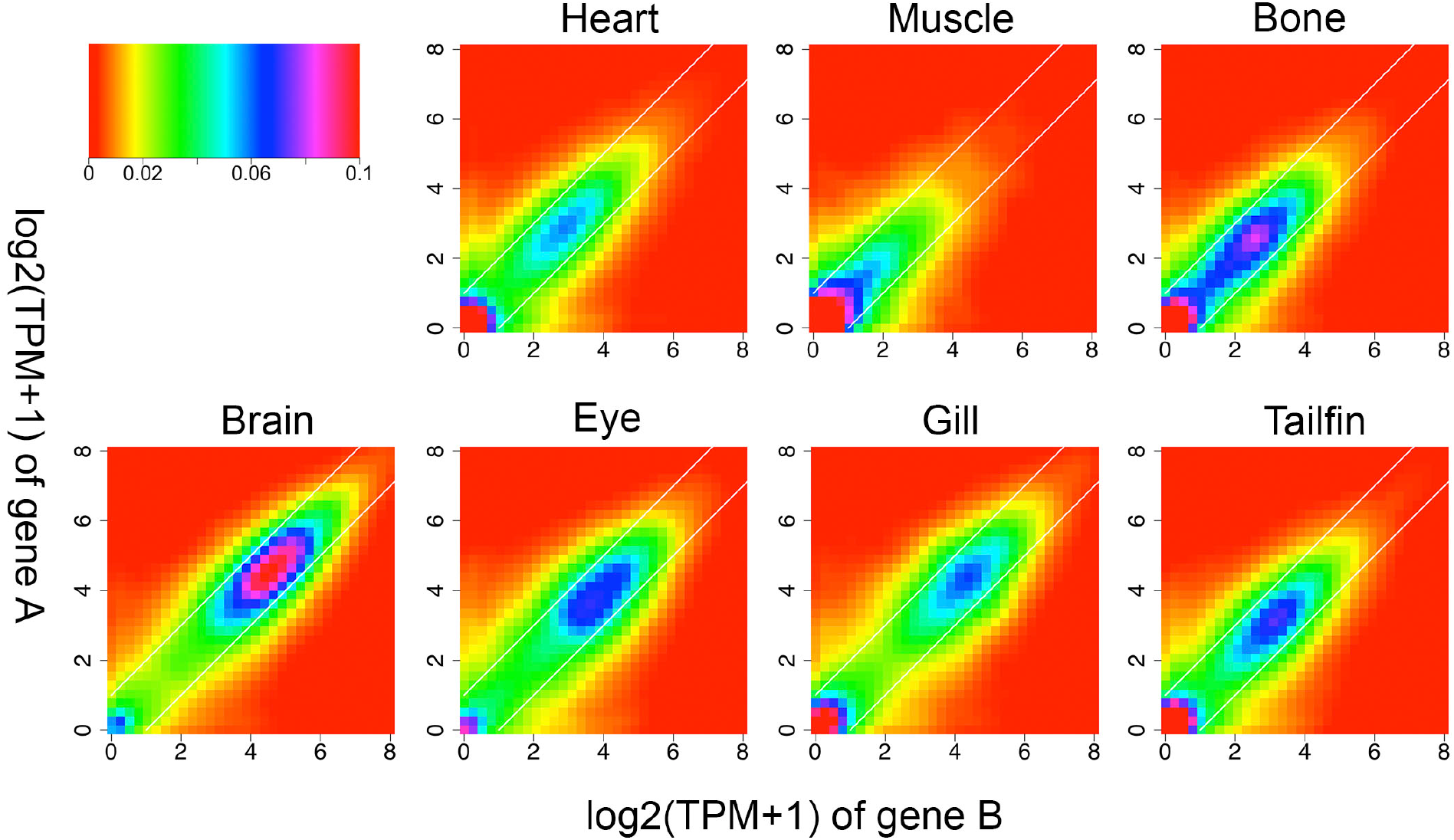
Expression of ohnolog gene pairs in seven tissues. Histogram is symmetrized. Color indicates percent of gene pairs. For each tissue, the TPM expression difference between most of gene pairs are less than 2-fold (i*.e.* between white lines).

**Supplemental Figure 7.**
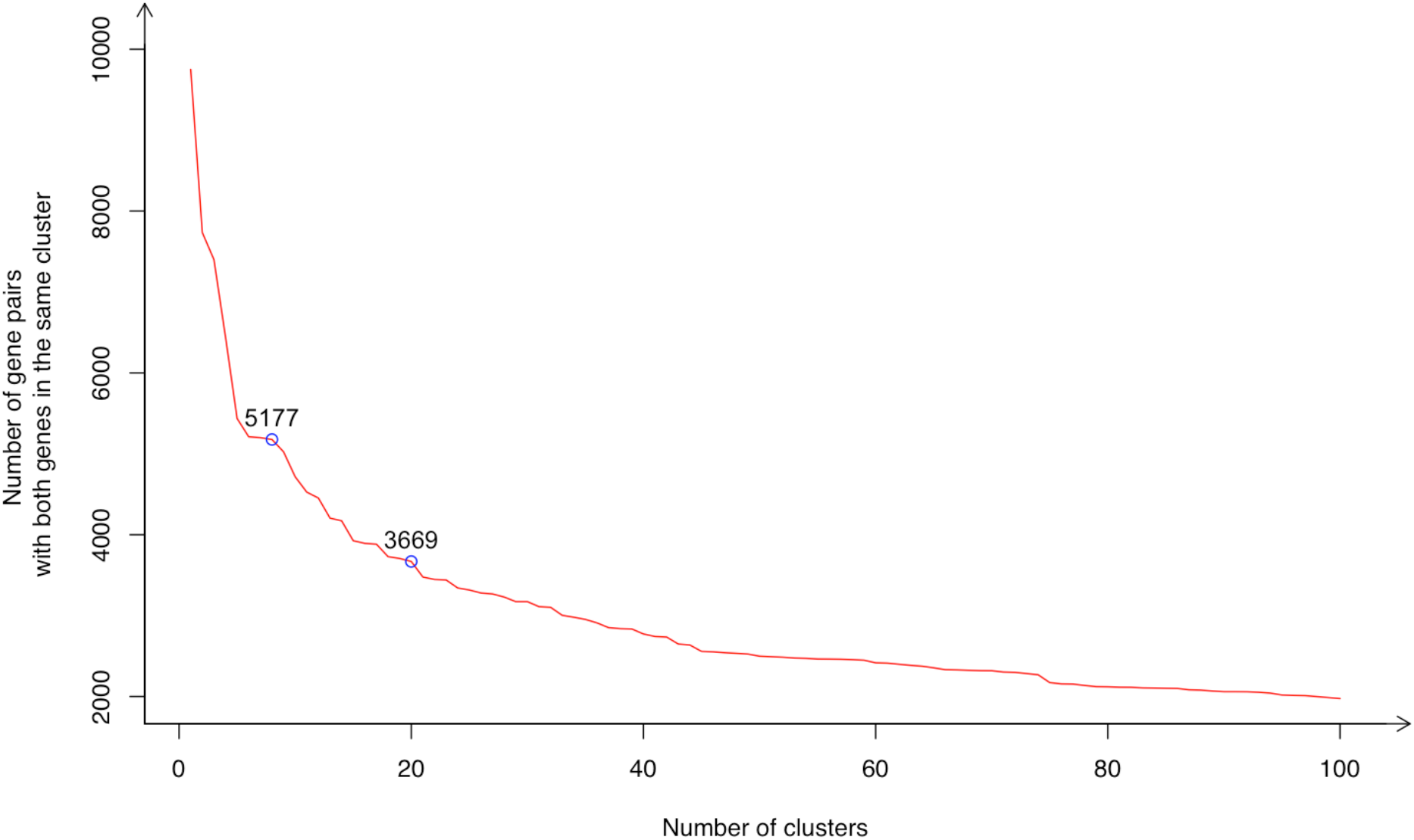
Total number of gene pairs with both ohnolog genes classified into the same expression cluster based on the number of clusters generated. Blue circles and the value shows the counts at 8 expression clusters and 20 clusters.

**Supplemental Figure 8.**
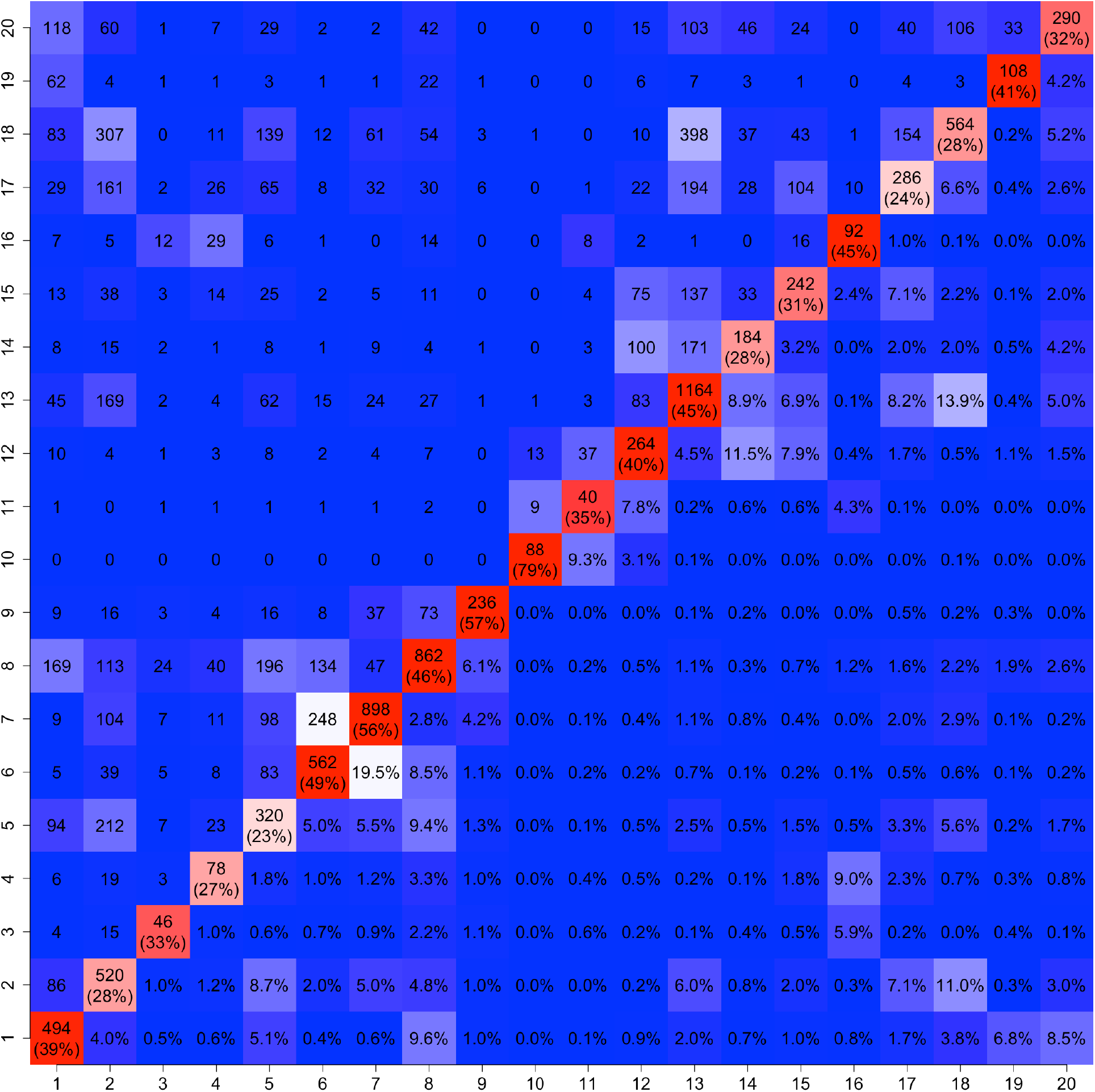
Number of ohnolog gene pairs in the same cluster (diagonal) or between each of the 20 clusters (upper triangle). The lower triangle shows the percentages. Blue-white-red Color indicates the percentage, from low to high.

**Supplemental Figure 15.**
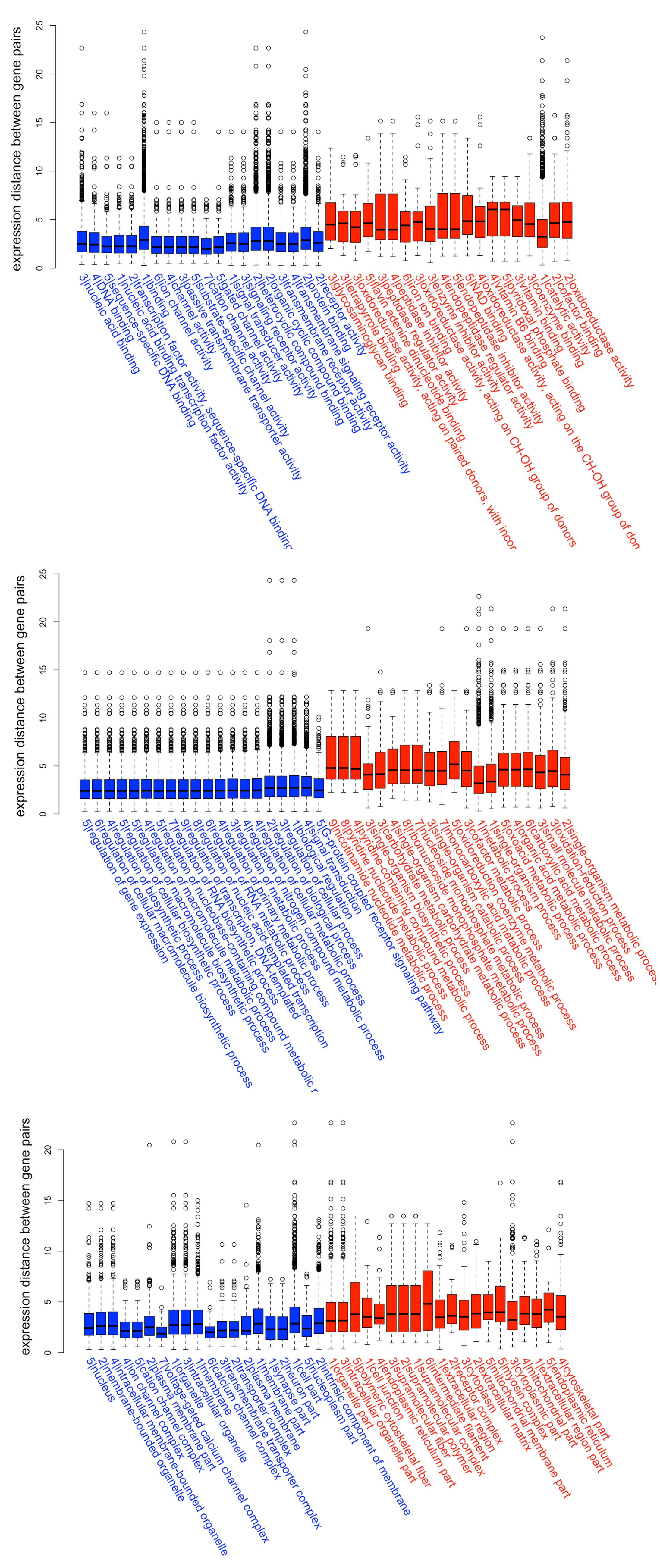
GO molecular function (**top**), biological process (**middle**) and cell component (**bottom**) with significantly low (top 20, blue) or high (top 20, red) expression distances between carp WGD ohnolog gene pairs (one side Wilcoxon rank sum test p<0.01).

